# Zero valent sulfur is the product of assimilatory sulfite reductase and a substrate of cysteine synthase

**DOI:** 10.1101/2024.12.13.628379

**Authors:** Qun Cao, Xuanyu Liu, Qingda Wang, Yongzhen Xia, Luying Xun, Huaiwei Liu

## Abstract

The long process of the assimilatory sulfate-reduction consists of sulfate reduction to H_2_S and then H_2_S assimilation for cysteine synthesis. However, recent studies indicate that both H_2_S and zero valent sulfur (S^0^) are produced during dissimilatory sulfate reduction process, which leads to the question whether S^0^ also is involved in assimilatory sulfate-reduction. Herein, phylogenetic analysis grouped assimilatory sulfite reductases into five clusters and we re-examined four representatives from two major clusters. These enzymes produced both S^0^ and H_2_S from sulfite. However, S^0^ was more like a direct product, while H_2_S was just a derivative of S^0^ via chemical reduction. Phylogenetic analysis of cysteine synthases grouped them into six clusters, and we re-examined six representatives from two major clusters. These enzymes used both glutathione persulfide (GSSH, a common intracellular compound containing a S^0^ atom) and H_2_S as substrates to synthesize cysteine, but preferred to use GSSH. These findings indicated that S^0^, other than H_2_S, is the key intermediate during the process of inorganic sulfur converting to organic sulfur.

**Importance:** Sulfate reduction is an important link in the global sulfur cycle. Assimilatory sulfate-reduction consists of sulfate-reduction to H_2_S and then H_2_S assimilation for cysteine synthesis. This concept was founded in 1950s and has not been challenged until today. In this study, we re-examined four representative assimilatory sulfite reductases (aSiRs) and found that these enzymes directly reduced sulfite to zero valent sulfur (S^0^). A small portion of produced S^0^ was reduced to H_2_S via aSiR-independent chemical reactions. Further, we re-examined six representative cysteine synthases and found that these enzymes used S^0^ derivative compounds (thiosulfate and glutathione persulfide) as substrates to synthesize cysteine. Thus, S^0^ is the key intermediate of assimilatory sulfate reduction process, other than H_2_S.

## Introduction

Sulfate reduction is a critical step of the sulfur biogeochemical cycle in which microorganisms play important roles. Depending on the fate of reduced sulfur atom, sulfate reduction process can be categorized into two types, dissimilatory sulfate reduction (DSR) and assimilatory sulfate reduction (ASR) (1, 2). DSR mainly happens in anaerobic environments where sulfate-reducing microorganisms (SRM) use sulfate as a terminal electron acceptor in the process of organic matter degradation. The canonical DSR process is thought to reduce sulfate to hydrogen sulfide (H_2_S) as the end product. However, zero valent sulfur (S^0^) has been widely detected in anaerobic conditions along with H_2_S (2). Recent studies indicated that dissimilatory sulfite reductases (dSiRs), specifically, DsrAB and DsrC generated S^0^ as an intermediate product (3–5).

Different from DSR, ASR is active in various environments where microorganisms use sulfate as a sulfur donor to synthesize organic compounds including cysteine, methionine, and glutathione. Sulfate is activated by an ATP sulfurylase (ATPS) to generate adenosine 5′-phosphosulfate (APS), which is directly reduced by an APS reductase (APSR) to generate sulfite. Alternatively, APS can be further phosphorylated to 3′-phosphoadenosine 5′-phosphosulfate (PAPS) by the APS kinase (APSK), which is then reduced to sulfite by a PAPS reductase (PAPSR) (6). Assimilatory sulfite reductases (aSiRs, some literatures used the name aSir, Asr, SiR and other names) finally reduced sulfite to H_2_S, which then reacts with O-acetyl-serine (OAS) to produce cysteine (6–9). Hence, H_2_S has been regarded as the key knot connecting sulfate-reduction and cysteine synthesis for over 60 years (10).

Most of early studies that focused on ASR ignored the presence and involvement of S^0^, mainly due to lack of related knowledge of S^0^ and its detection methods (11–15). It is possible that S^0^ also is present in ASR process because, first, recent studies indicated that S^0^ is widely present in all types of cells. It is not stable and can easily react with H_2_S, small cellular thiols, such as glutathione (GSH), and sulfite to form hydrogen polysulfides (HS_n_H, n≥2), glutathione polysulfides (GS_n_H, n≥2), and thiosulfate (S_2_O_3_ ^2-^) (16–20). Second, phylogenetic sequence analysis indicates that aSiRs diverged from the same ancestral gene as dSiRs (21–23). Therefore, aSiRs may share similar catalytic mechanisms as dSiRs and also product S^0^, but no related study has been reported as so far.

In recent years, a number of H_2_S producing enzymes have been redefined as S^0^ producing enzymes thanks to the development of S^0^ analysis methods. For instance, 3-mercaptopyruvate sulfurtransferase (3-MST), which was reported to cleave the C–S bond of 3-mercaptopyruvate to produce H_2_S. In 2015, Kimura *et al*., discovered that the released sulfur atom actually existed as HS_n_H (24). Cystathionine β-synthetase (CBS) and cystathionine γ-lyase (CSE) both produced S^0^ from cystine (25, 26). Methanethiol oxidase (MTO) catalyzed methanethiol degradation to produce S^0^ other than previously reported H_2_S (27). From sulfur cycle viewpoint, these findings indicate that when organic sulfur is converted to inorganic sulfur by biological systems, S^0^ is the direct product.

Previous studies indicated that H_2_S is the main substrate of cysteine synthases (28), but a few studies found that some microbial cysteine synthases also can use thiosulfate as an alternative substrate to produce cysteine, which is less efficient probably due to thiosulfate is weaker than H_2_S in aspect of nucleophilicity and thiosulfate forms a disulfide bond intermediate with OAS before generates cysteine (29, 30). In our previous study, we discovered that the *Schizosaccharomyces pombe* cysteine synthase Cys11 uses glutathione persulfide (GSSH) as a substrate to produce cysteine (31). Moreover, for Cys11, the *K_m_* value of GSSH is ∼400 μM, 2-fold lower than that of H_2_S (∼800 μM). Knocking out the sulfide:quinone oxidoreductase (Sqr), which converts H_2_S to S^0^ in mitochondria, leads to cysteine-deficient phenotypes. Therefore, we proposed that S^0^ might be the *defacto* substrate of Cys11 inside of *S. pombe* cells. Whether Cys11 is an exception or it represents a common characteristic of microbial cysteine synthases needs further investigation.

In this study, we performed phylogenetic analysis of sulfite reductases and cysteine synthases. Representative enzymes were purified and their activities were examined. We found that all examined assimilatory sulfite reductases (aSiRs) produced more S^0^ than H_2_S from sulfite. H_2_S was a derivative product of S^0^. Microorganisms overexpressing aSiRs accumulated more S^0^ than H_2_S from sulfite reduction. All representative cysteine synthases can use GSSH as a substrate to synthesize cysteine and most of them preferred GSSH to H_2_S. These findings suggest that as in the H_2_S oxidation pathway, S^0^ also is the intermediate in the ASR pathway. S^0^ is more like the key knot connecting sulfate reduction and cysteine synthesis other than H_2_S.

## Results

### Methylene blue method cannot distinguish between hydrogen sulfide and hydrogen persulfide

Methylene blue method was commonly used in early studies of sulfite reductases. The mechanism is that two molecules of *N,N*-dimethyl-p-phenylenediamine (DMPD) and one H_2_S molecule are oxidized by ferric ion (Fe^3+^) to form a methylene blue dye, which has a maximal absorbance at around 670 nm (A_670_) (Fig. 1). The A_670_ value is linearly correlated with amount of H_2_S in the presence of excessive Fe^3+^/DMDP (32, 33). We found that Fe^3+^/DMPD not only reacted with H_2_S, but also with hydrogen persulfide (HSSH). Furthermore, the latter generated higher A_670_ values than the former at the same doses (Fig. 1A). LC-ESI-MS analysis indicated that when H_2_S reacted with Fe^3+^/DMDP, not only methylene blue compound (**a**) was produced, another compound whose structure is similar as **a** but contains an extra sulfur atom (**b**) was also produced (Fig. S1). We proposed that **a** and **b** were generated from reactions 2 and 3, respectively (Fig. 1B). When HSSH reacted with Fe^3+^/DMDP, **a** and **b** also were produced. The detailed mechanism was not investigated, but could be because HSSH is inherently unstable and easily breaks into H_2_S and S^0^ in aqueous solution, or HSSH could react with Fe^3+^/DMDP directly (reactions 4 and 5).

**FIG 1.**
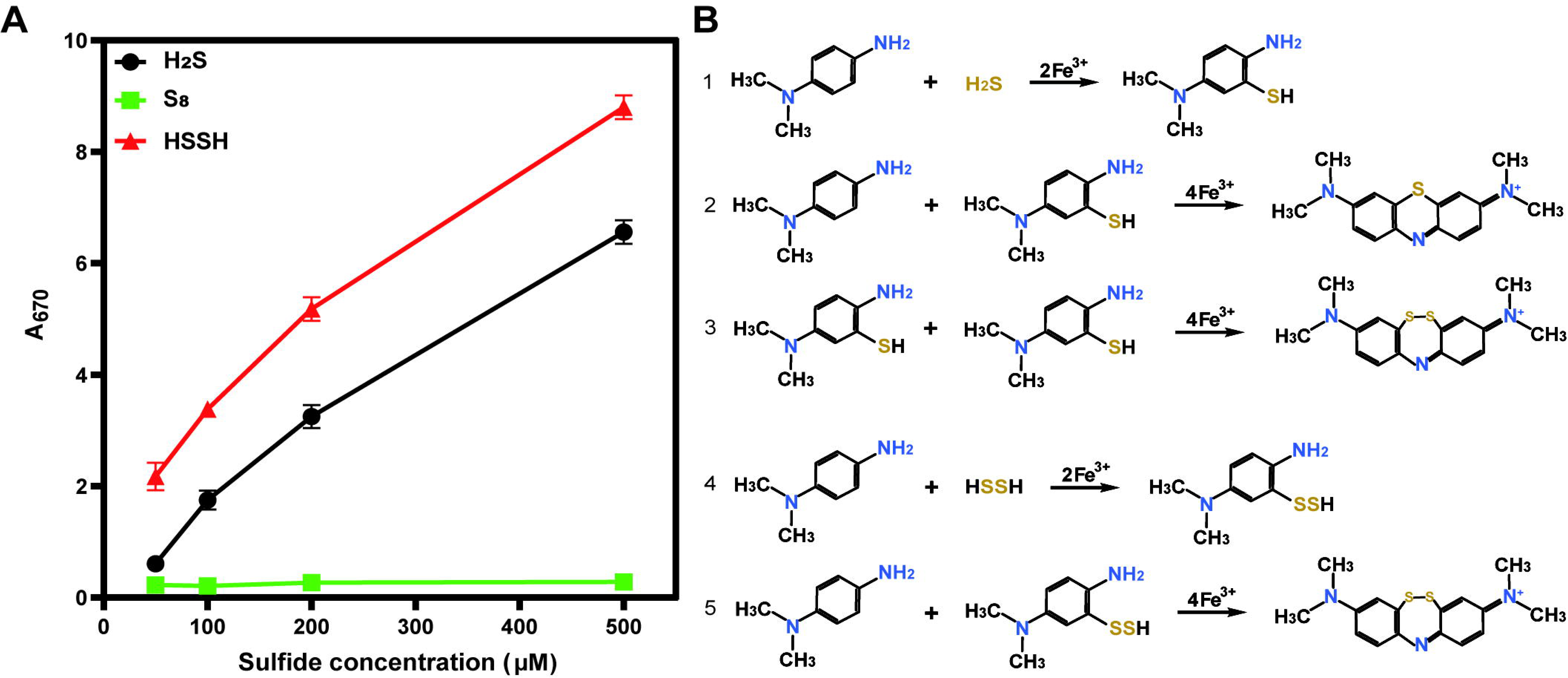
Application of methylene blue method in H_2_S and S^0^ quantification. (A) HSSH and H_2_S can react with methylene blue reagents to generate thiaazaheterocyclic dyes but S_8_ cannot. Data are from three replicated experiments (n=3) and presented as average ± s.d; (B) Mechanisms of how HSSH and H_2_S react with methylene blue reagents. Reactions 1 and 2 are from reference (24). Other reactions are proposed by authors.

Elemental sulfur (S_8_) did not generate any methylene blue dye, suggesting that only nucleophilic polysulfide (polysulfide with terminal hydrogen atoms) can react with Fe^3+^/DMDP to produce S-containing heterocyclic dyes (Fig. 1B).

### Representative aSiRs produced more S^0^ than H_2_S from sulfite in vitro

A total of 457,826 protein sequences are denoted as sulfite reductase in GenBank (update to August 29, 2024). These sequences were downloaded and subjected to redundancy analysis. After removing redundant sequences, 4,378 sulfite reductases were left. Their characteristic sequences were identified (PF01077 and PF03460) and using the presence or absence of the identified characteristic sequence as a screening criterion, 289 sulfite reductases were finally extracted out. Origins of these enzymes encompass bacteria, archaea, fungi, and plants. Phylogenetic analysis of these enzymes indicated that 289 enzymes were categorized into five major clusters with clusters 3 and 4 were two largest ones (Fig. 2 and Table S1).

**FIG 2.**
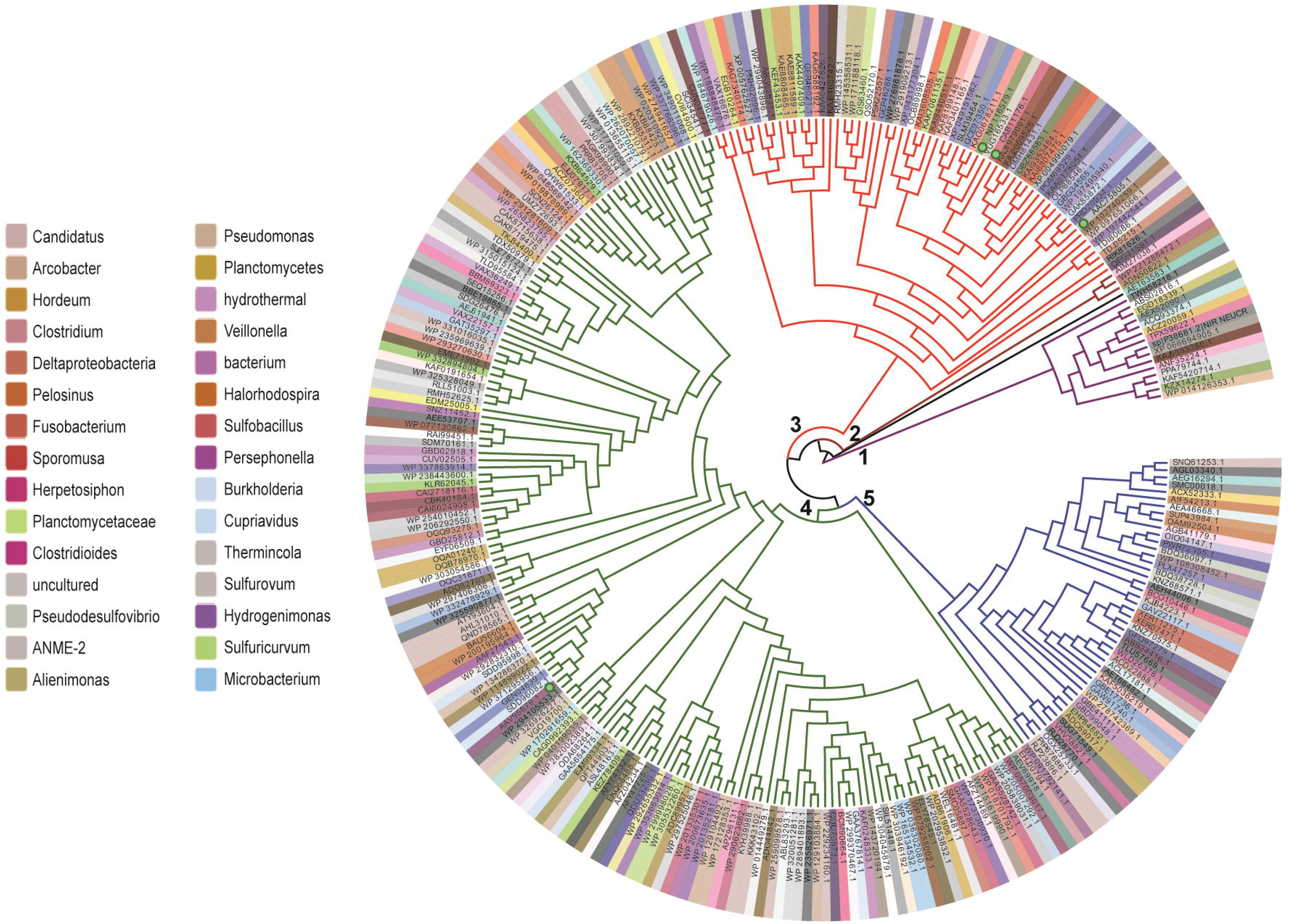
Phylogenetic analysis of 289 sulfite reductases. Only part origins of the enzymes were shown. Details of these enzymes and their origins are shown in Table S1.

We chose four aSiRs as representatives for further examination: *E. coli* CysIJ (AAC75805.1), *S. cerevisiae* MET5&MET10 (NP_ 116579.1), and *S. pombe* Sir1 (CAB11176.1). They all belong to cluster 3; *Ruegeria pomeroyi* DSS-3 SPO2634 (AAV95879.1) belongs to cluster 4. Among them, CysIJ contains two subunits (Cys I and Cys J) and has been intensively studied as a model aSiR in 1960s (10). Others are from widely used model microorganisms. *S. cerevisiae* MET5&MET10 also contains two subunits, while *S. pombe* Sir1 and *R. pomeroyi* DSS-3 SPO2634 are single subunit enzymes. These subunits/enzymes were fused with an N-terminal His-Tag, expressed in *E. coli* BL21(DE3), and purified using a nickel column (Fig. S2). During cell disruption and protein purification processes, no dithiothreitol (DTT) or other reductants was added. Freshly purified enzymes were immediately subjected to activity analysis.

We mixed purified CysI and CysJ to obtain the CysIJ complex. Likewise, the MET5&MET10 complex was obtained. CysIJ, MET5&MET10, and Sir1 were NADPH dependent enzymes, we mixed them with NADPH, hemin, and sulfite. Since SPO2634 was a ferredoxin dependent enzyme, we mixed it with ferredoxin (Fd), ferredoxin-NADP+ oxidoreductase (FNR), NADPH, and sulfite. Fd and FNR encoding genes were amplified from genomic DNA of *R. pomeroyi* DSS-3. We expressed them in *E. coli* BL21(DE3) and purified these two proteins for this experiment. To analyze the produced sulfur compound, we used a LC-MS dependent method instead of the methylene blue method. The enzymatic products were derivatized with monobromobimane (mBBr) and analyzed using LC-ESI-MS. For the reaction catalyzed by CysIJ, H_2_S derivatives mB-SH and mB-S-mB were detected. Signal intensity of the former was 1.9×10^6^ and the latter was 4.6×10^5^ (Fig. 3A, Fig. S3A and S3B). Surprisingly, derivative of HSSH (mB-SSH) also was detected and its signal intensity was 3.6×10^6^ (Fig. 3A, Fig. S3C). Previous studies indicated that when derivatizing H_2_S and HSSH with mBBr, dominant products were mB-S-mB and mB-SS-mB, respectively (34). Here, we detected two derivatives of H_2_S, mB-S-mB and mB-SH (partial derivatization product), and one derivative of HSSH, mB-SSH (partial derivatization product). The most probable reason is aSiRs and sulfite also react with mBBr, which disturbs the reaction between H_2_S/HSSH and mBBr, and leads to partial derivatization. Considering mB-SH and mB-S-mB have similar fluorescence and structure, we summed their signal intensities (roughly representing H_2_S amount) and compared it with signal intensity of mB-SSH (roughly representing HSSH amount). The latter was higher than that of the former. Similarly, signal intensity of H_2_S derivatives (mB-SH + mB-S-mB) from MET5&MET10 was 2.4×10^6^; whereas signal intensity of HSSH derivative (mB-SSH) was 3.2×10^6^ (Fig. 3B). Signal intensities of H_2_S derivatives from SPO2634 and Sir1 also were lower than that of HSSH derivative (Fig. 3C and 3D).

**FIG 3.**
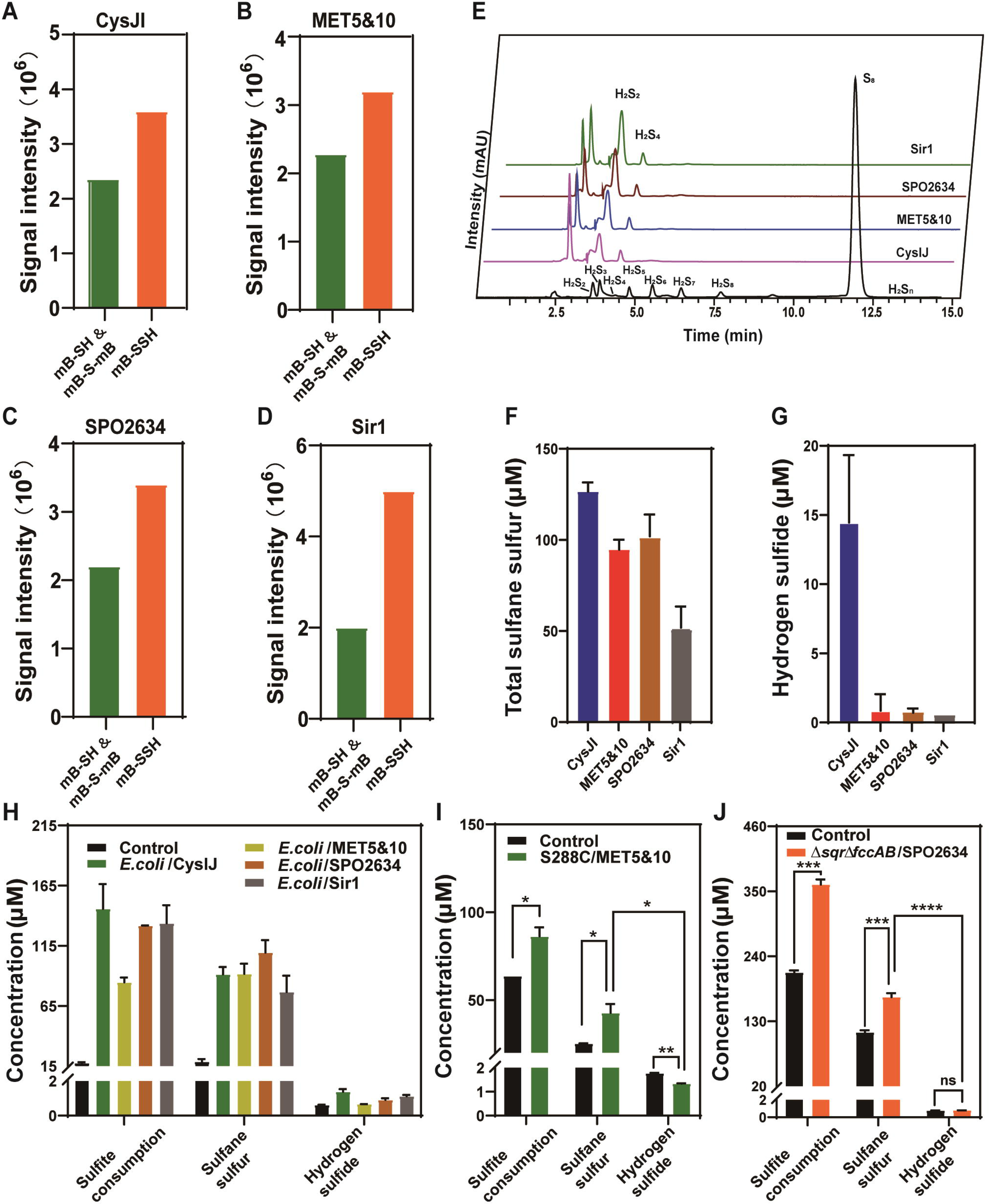
Analysis of the products generated from sulfite reductase-catalyzed reactions. Purified CysIJ (A), MET5&10 (B), and SPO2634 (C) were mixed with NADPH andsulfite; purified Sir1 (D) was mixed with ferredoxin, ferredoxin-NADP+ oxidoreductase, NADPH, and sulfite. The reactions were conducted at 25°C for 10 min. Products were derivatized by mBBr. After mBBr derivatization, H_2_S became mB-SH and mB-S-mB, and HSSH became mB-SSH. The derivatives were detected by LC-ESI-MS. MS signal intensities of H_2_S (mB-SH + mB-S-mB) and HSSH (mB-SSH) were shown. MS spectra of these chemicals were shown in Fig. S3; (E) The enzyme-produced H_2_S_n_ was derivatized by methyl trifluoromethanesulfonate and analyzed using HPLC with UV detection. H_2_S_n_ mixture was used as the standard; (F) The enzyme-produced H_2_S_n_ was quantified using cyanide method and shown as total S^0^; (G) The enzyme-produced H_2_S was quantified by HPLC; (H) Sulfite consumption, S^0^ production, and H_2_S production from *E. coli* MG1655 strains overexpressing sulfite reductases; (I) Sulfite consumption, S^0^ production, and H_2_S production from *S. cerevisiae* S288C overexpressing MET5&10; (J) Sulfite consumption, S^0^ production, and H_2_S production from *R. pomeroyi* DSS-3Δ*sqr*Δ*fccAB* overexpressing SPO2634. Controls are strains without sulfite reductase overexpression. For F–J, data are from three replicated experiments (n=3) and presented as average ± s.d. T-tests were performed to calculate the p-values. Asterisks indicate statistically significant difference (* p < 0.05, ** p < 0.01, ***p < 0.001, ****p < 0.0001).

A previous report indicated that hemin itself can catalyze sulfite reduction (35). The authors used methyl viologen (MeV) as electron source. By monitoring the decrease in absorbance at 600 nm from the MeV radical cation, they calculated the reaction kinetics and concluded that free siroheme was a more efficient catalyst than dSiRs. No product analysis was performed in that study but the authors proposed that trithiolate was a product. Here, for CysIJ, MET5&MET10, and Sir1 catalyzing reactions, hemin was added as a cofactor. To find out whether hemin itself can reduce sulfite to H_2_S or HSSH, we mixed hemin with sulfite. The reaction conditions were the same except that no sulfite enzyme was added. LC-ESI-MS analysis demonstrated that no H_2_S or HSSH was detected.

To further confirm the S^0^ products of these enzymes, we analyzed the products with the trifluoromethanesulfonate derivatization-based method, which can discriminate polysulfides with different length of chains (H_2_S_n_, n=2-8) (36). The reaction mixtures were reacted with methyl trifluoromethanesulfonate to convert hydrogen polysulfide (H_2_S_n_) to dimethylpolysulfide. Results showed that HSSH (H_2_S_2_) was the main product of these enzymes while HSSSSH (H_2_S_4_) was the minor one (Fig. 3E). We then quantified the produced total sulfane sulfur (representing amount of S^0^ atoms) with a cyanide method and compared it to the amount of produced H_2_S. Results showed that CysIJ, MET5&MET10, and SPO2634 produced 80∼130 μM S^0^, Sir was less efficient than them and produced about 50 μM S^0^ from 150 μM sulfite. In comparison, CysIJ produced about 15 μM H_2_S and the other three aSiRs produced only 1∼5 μM H_2_S (Fig. 3F and 3G). These results indicated that all of the four aSiRs produced more S^0^ than H_2_S.

For the presence of HS_n_H in the products, a possible explanation is that aSiRs firstly produce H_2_S from sulfite, and HS_n_H are generated from autonomous oxidation of H_2_S (by dissolved oxygen). However, according to our experience, the process of H_2_S autonomous oxidation is slow, which usually takes hours without the help of related enzymes or strong oxidants (such as Fe^3+^ and H_2_O_2_). For confirmation, we mixed H_2_S with reaction buffer containing no aSiR. After 10 min incubation at 25°C, no HS_n_H was produced, suggesting that HS_n_H was not produced from autonomous oxidation of H_2_S. As the control, we also mixed HSSH with reaction buffer containing no aSiR and other reagents. Other conditions were the same as that of enzymatic reaction. H_2_S production was detected and its amount was about 1/10 of HSSH. Considering that HSSH is easily reduced to H_2_S when reductant is present, we also mixed HSSH, reaction buffer, and NADPH. In this case, H_2_S production was significantly increased to about 1/2 of HSSH (Fig. S4).

Together, the above results suggested that under the tested conditions, S^0^ was the direct product of all four aSiRs. As showing in equation 1, aSiR gets four electrons from NADPH and passes them to sulfite. Sulfite is reduced to S^0^. The produced S^0^ atoms either attach to the enzyme and exist as protein-S_n_H, or exist as free S_n_ (equation 2). In either case, S_n_ can get more electrons and become water soluble HS_n_H (equation 3). When there is abundant NADPH present, HS_n_H is easily reduced to H_2_S (equation 4). Since the H_2_S generation reaction is aSiR independent, H_2_S is more like a derivative product of ASR process, not the direct product.

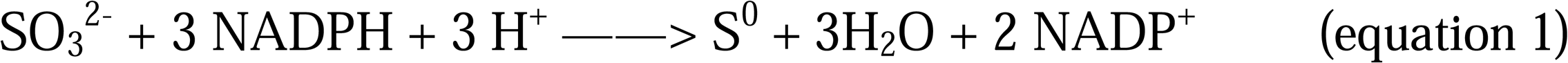

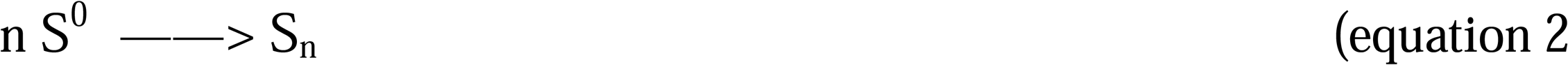

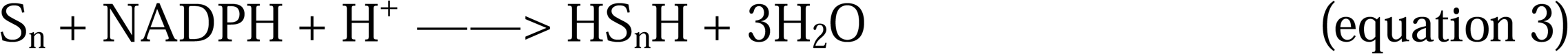

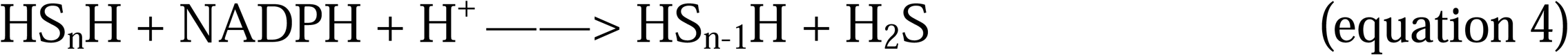

### Representative aSiRs produced more S^0^ than H_2_S in vivo

Since *E. coli* MG1655 contains no H_2_S oxidation enzyme, testing the products of aSiRs in it is convenient. We expressed the four aSiRs (using the pBBR1MCS5 plasmid as a vector) in *E. coli* MG1655. *E. coli* MG1655 with the pBBR1MCS5 empty plasmid was used as a control. The cell suspension (10 ml, OD_600_=3) was mixed with 0.5 mM sulfite in PBS buffer (pH 7.4, 50 mM) and the mixture was incubated at 37□ with shaking. After 1 h incubation, consumed sulfite and generated products were quantified. We observed that compared with the control, all aSiR expression strains consumed more sulfite. More importantly, they all produced more total S^0^ than H_2_S from the consumed sulfite (Fig. 3H).

We also constructed a *S. cerevisiae* S288C strain (that contains no *sqr*) with MET5&MET10 overexpression and a *R. pomeroyi* DSS-3 Δ*sqr*Δ*fccAB* strain with SPO2634 overexpression (Since *sqr* and *fccAB* encode H_2_S oxidation enzymes, deleting these two genes prevent the oxidation of H_2_S to S^0^). Cell suspensions of these strains were mixed with sulfite, and the products were analyzed. Similarly, they also produced more S^0^ than H_2_S (Fig. 3I and 3J). These results demonstrated that the tested aSiRs tended to produce S^0^ other than H_2_S *in vivo*.

### Representative cysteine synthases utilized S^0^ to produce cysteine

A total of 499,263 protein sequences are denoted as cysteine synthase in GenBank (update to August 29, 2024). The similar screening process was performed and 735 cysteine synthases were finally extracted out. Origins of these enzymes encompass bacteria, archaea, fungi, and plants. Phylogenetic analysis of these enzymes indicated that they were categorized into six major clusters with clusters 4 and 5 were two largest ones (Fig. 4, Table S2).

**FIG 4.**
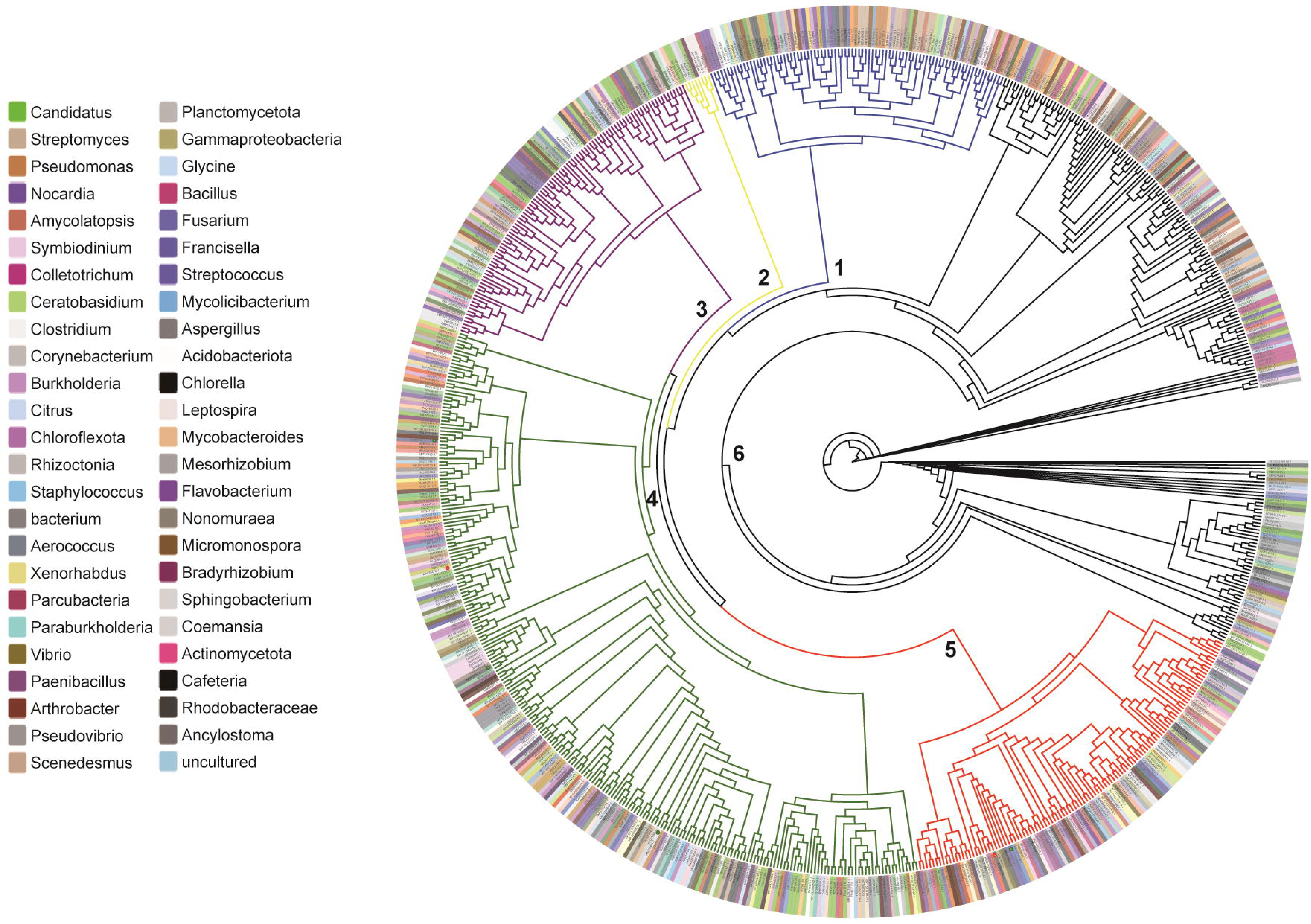
Phylogenetic analysis of 735 cysteine synthases. Only part origins of the enzymes were shown. Details of these enzymes and their origins are shown in Table S2.

We selected 5 microbial cysteine synthases for further study: CysK and CysM from *E. coli* MG1655 (EcCysK and EcCysM), CysK from *R. pomeroyi* DSS-3 (RpCysK), SiRe_ 1641 from *Sulfolobus islandicus* REY15A, a model archaeon that belongs to Crenarchaeota. They all belong to cluster 4. CysM from *R. pomeroyi* DSS-3 (RpCysM) belongs to cluster 5. We expressed their encoding genes in *E. coli* BL21(DE3). These enzymes (fused with a His-tag at N-terminus) were purified using a nickel column (Fig. S5). Freshly purified enzymes were subjected to activity analysis immediately.

A catalytic mechanism of cysteine synthase has been reported. OAS first reacted with PLP to form the α-aminoacrylate intermediate containing a C=C bond, which was electrophilic. Subsequently, nucleophilic H_2_S attacked the C=C bond of α-aminoacrylate to generate cysteine (Fig. 5) (37). We mixed purified cysteine synthases with H_2_S/GSSH/Na_2_S_2_O_3_, OAS, and pyridoxal 5-phosphate (PLP) in PBS buffer (50 mM, pH 7.4) and incubated the mixture at 30°C for 5 min. LC-ESI-MS method was used to analyze the products.

**FIG 5.**
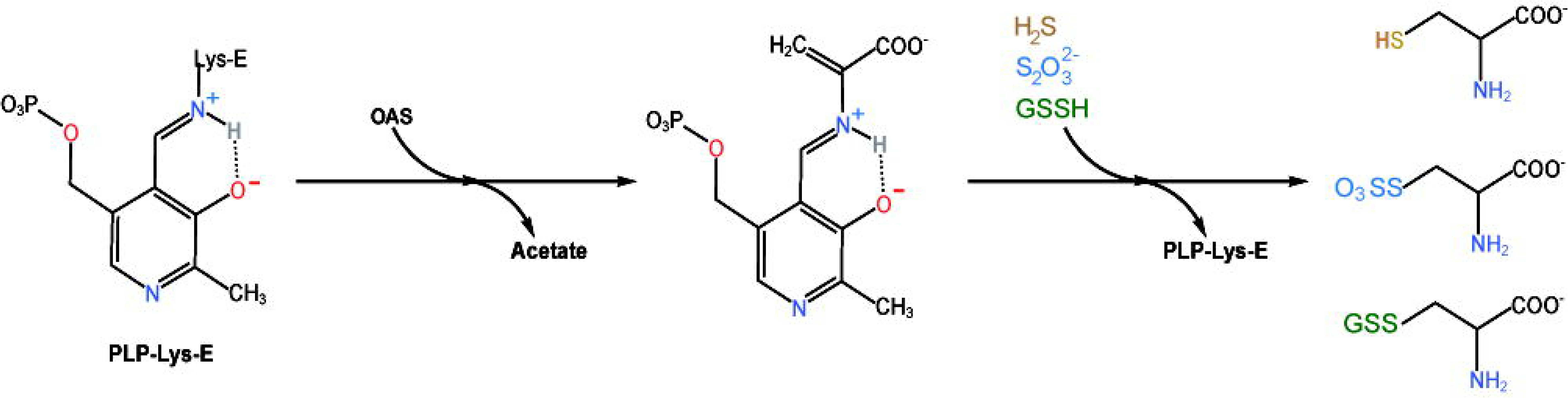
Pathway of cysteine synthesis using different sulfur donors.

From the catalyzing mechanism perspective, GSSH and Na_2_S_2_O_3_ react with OAS will generate GS-cysteine and sulfo-cysteine intermediates, respectively (equations 5-6). We detected these two intermediates from reactions catalyzed by all five enzymes (Fig. S6-S11). After treating the intermediates with tris-(2-carboxyethyl)-phosphine (TCEP), L-cysteine was detected (equations 7-8). These enzymes also used H_2_S as substrate to produce L-cysteine (equation 9).

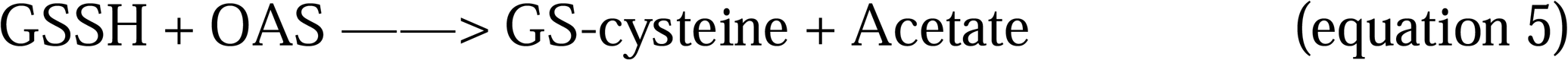

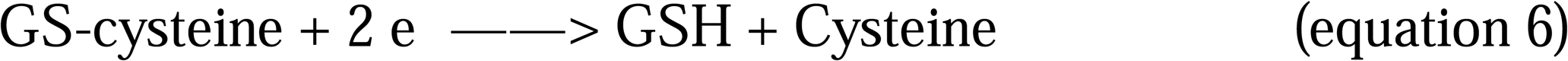

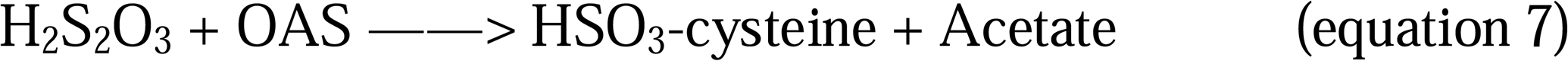

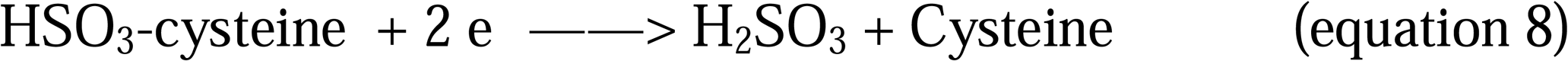

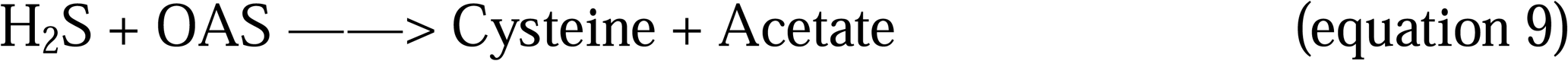

We then assayed the kinetics of all five cysteine synthases (Fig. 6, Fig. S12-S14). The obtained data were compared with the reported data of *S. pombe* cysteine synthase Cys11 (31). For all 6 enzymes, the *K_m_*values for GSSH were between 202∼426 μM, *K_m_* values for H_2_S were between 157∼3192 μM, and *K_m_* values for Na_2_S_2_O_3_ were between 408∼3774 μM. The *V_ma_*_x_ values for GSSH were between 0.125∼0.280 mM·min^-1^, *V_ma_*_x_ values for H_2_S were between 0.139∼0.581 mM·min^-1^, and *V_ma_*_x_ values for Na_2_S_2_O_3_ were between 0.021∼0.173 mM·min^-1^. The *K_cat_* values for GSSH were between 2.083∼9.778 s^-1^, *K_cat_* values for H_2_S were between 3.127∼13.833 s^-1^, and *K_cat_* values for Na_2_S_2_O_3_ were between 0.495∼2.454 s^-1^. These results indicated that among GSSH, H_2_S, and Na_2_S_2_O_3_, Na_2_S_2_O_3_ was the most unpreferred substrate. Most cysteine synthases showed higher affinity to GSSH than to H_2_S.

**FIG 6.**
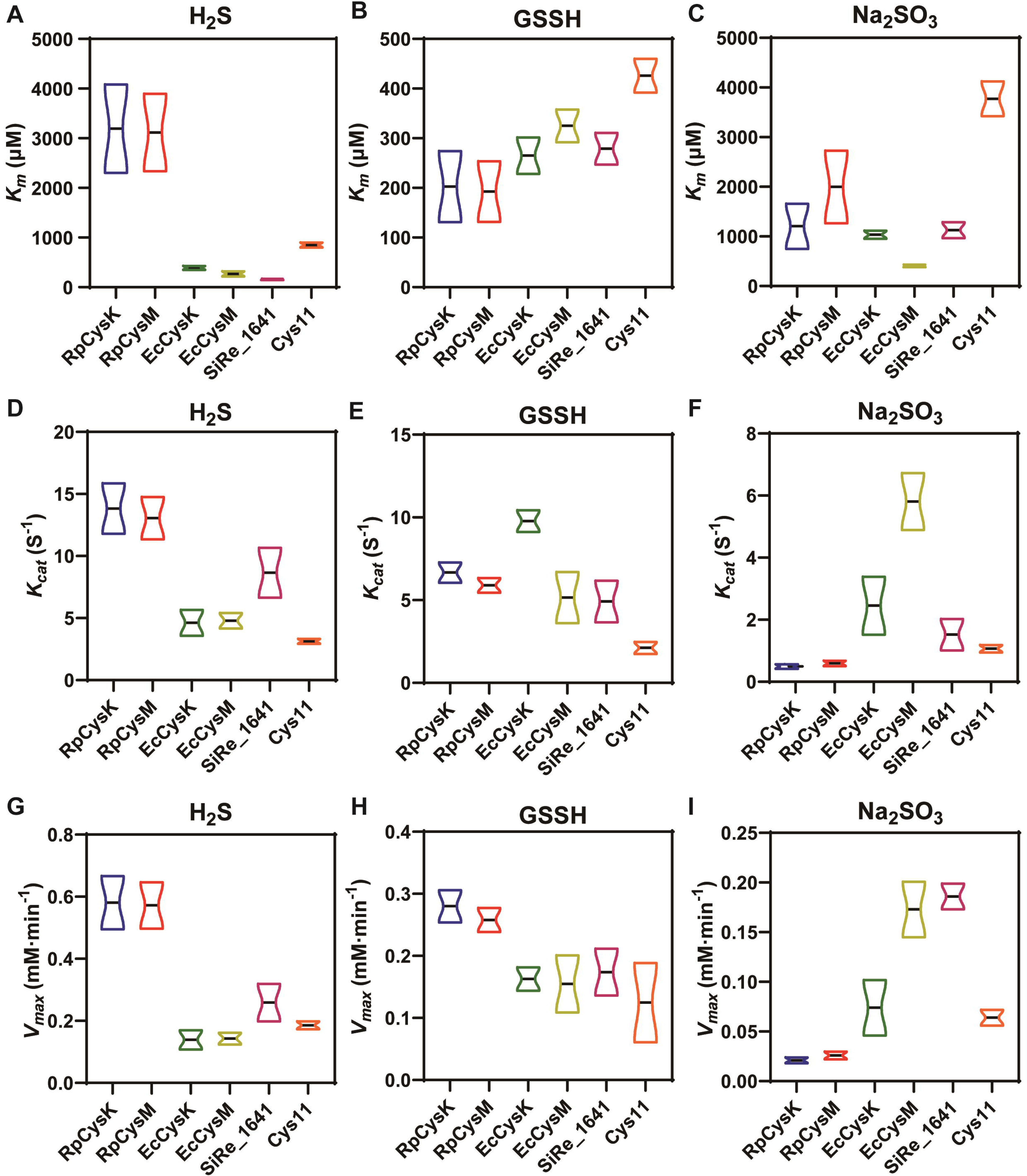
Kinetic assays of cysteine synthases with different substrates. Details of the data are shown in Table S3 and Fig. S12–S14.

## Discussion

Inorganic sulfur (sulfate) becomes organic sulfur via the assimilatory sulfate reduction and cysteine synthesis pathways. The concept that H_2_S is a key intermediate of this process is founded in 1960s when S^0^ analysis methods were short and knowledge of biological S^0^ were scarce (10). In this study, we re-examined the product of aSiRs with modern methods that can distinguish S^0^ containing compounds from H_2_S. We found that all tested aSiRs produced both S^0^ and H_2_S from sulfite. S^0^ was more like the direct product of aSiRs while H_2_S was its derivative. We also re-examined model cysteine synthases and discovered that all tested cysteine synthases can use S^0^ (S_2_O_3_^2-^ and GSSH) as alternative substrates to produce cysteine.

From chemistry viewpoint, biological S^0^ compounds are better intermediates than H_2_S. First, the p*K_a_* 1 of H_2_S is 6.8 and gasification temperature is 34.3°C, so it easily releases out of cells under normal physiological condition. In comparison, the p*K_a_* of GSSH is 5∼6, and therefore, it mainly exist in deprotonated form inside cells (38). Thiosulfate is even more stable than GSSH. Second, H_2_S is toxic to the respiration system, its intracellular concentration is limited to 10∼30 nM level (33), which was far below the *K_m_* value of cysteine synthases. In contrast, intracellular S^0^ chemicals are high to 100∼400 μM (39, 40), which was near the *K_m_* values of cysteine synthases. Third, biological S^0^ compounds have distinctive activities. For instance, GSSH is more nucleophilic than GSH because of the α-effect and hence displays high reactivity toward electrophiles such as reactive oxygen species (ROS) and toxic metal cations (41). In consistent, we found that cysteine synthases preferred GSSH the most. It is noteworthy that GSSH leads to the production of GS-cysteine intermediate which contains a disulfide bond. We used TCEP to cut this disulfide bond to release cysteine *in vitro*. It is still unknown how this disulfide bond is cleaved in the cell, small reducing molecules like GSH might be responsible for this reaction. In addition, we also found that microorganisms accumulated high concentration of intracellular S^0^ during the fast growth phase (cysteine synthesis process is active in fast growth phase), but reduced its level quickly after that (40). Combining the above evidence, we proposed that S^0^ is the *de facto* substrate of cysteine synthases *in vivo*. S^0^ probably is a key knot connecting sulfate reduction and cysteine synthesis in microorganisms. H_2_S is more like an efflux of intracellular S^0^ (Fig. 7).

**FIG 7.**
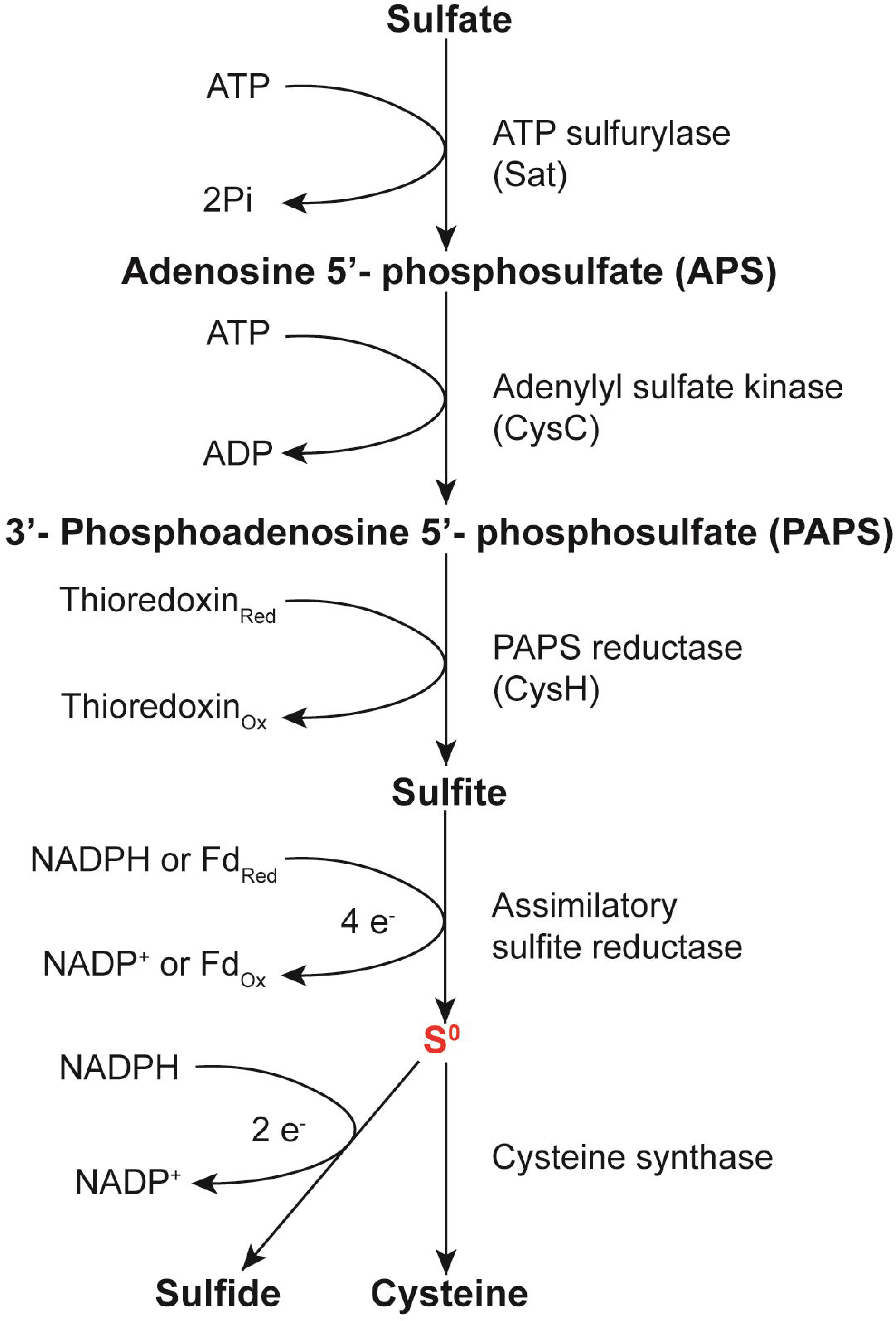
Process of the assimilatory sulfate reduction. Zero valent sulfur (S^0^) is the key intermediate linking sulfate reduction and cysteine synthesis and H_2_S is a side-product.

Recent studies indicated that S^0^ is an intermediate product in DSR process (4). A proposed mechanism is, first, DsrAB complex reduces sulfite to S^0^, which then is transferred to DsrC in the form of DsrC-SSH or DsrC trisulfide (3, 5). Second, DsrC-SSH or DsrC trisulfide receives electrons from the Dsr membrane complex and reduce the bound S^0^ to H_2_S. Therefore, DsrC is an important S^0^ carrier in this process. Some bacteria contain only DsrABC and no Dsr membrane complex. These bacteria may only produce S^0^ during sulfate reduction and DsrC may transfer its S^0^ to other acceptors in the cell. Using the *Archaeoglobus fulgidus* DsrC as seed and 35% homology as criteria, we searched for the DsrC homologs in genomes of *E. coli*, *S. cerevisiae*, *S. pombe*, and *R. pomeroyi*, no DsrC homologue was found. Probably because DsrC mainly exists in anaerobic bacteria but the four strains are all aerobic/facultative bacteria. No evidence indicates that DsrC is involved in ASR process as so far. In conclusion, we found that like in DSR process, S^0^ presents in ASR process as a direct product of aSiRs, probably because both aSiRs and dSiRs diverge from a common ancestor.

## Materials and methods

### Strains and cultivation conditions

*R. pomeroyi* DSS-3 is a gift from Prof. Yuzhong Zhang of Ocean University of China. *R. pomeroyi* DSS-3 derivative Δ*sqr*Δ*fccAB* were constructed in a previous work(29). *S. islandicus* REY15A is a gift from Prof. Yulong Shen of Shandong University. *S. pombe* YHL6381 is a gift from Prof. Huang Ying. *E. coli* strains used for plasmid construction and protein expression, and plasmids constructed in this study are all listed in Table S4. *E. coli* strains were cultured in LB medium. *R. pomeroyi* DSS-3 strains were cultured in 1/2 YTSS medium. *S. cerevisiae* S288C strains were cultured in YPD or SD medium. *S. pombe* YHL6381 strains were cultured in YES or SD medium. *S. islandicus* REY15A were grown in MTSV medium [mineral salt supplemented with 0.2% w/v (weight/volume) tryptone (T), 0.2% (w/v) sucrose (S), mixed solution of vitamins (V)] or MTSVU medium [MTSV supplemented with 0.001% (w/v) uracil (U)]. For cultivation, *E. coli* strains were cultured at 37□ with shaking (220 rpm). *R. pomeroyi* DSS-3, *S. cerevisiae* S288C and *S. pombe* YHL6381 strains were cultivated at 30□ with shaking (220 rpm). *S. islandicus* REY15A strains were cultivated at 75□ with shaking (220 rpm).

### Reagents

Sodium hydrosulfide (NaHS), sodium sulfide (Na_2_ S), cysteine, reduced glutathione (GSH), sulfite and thiosulfate were purchased from Sigma-Aldrich (Saint Louis, MO). S8 were purchased from TCI Company (Shanghai, China). HSSH was prepared following the protocol of Xin *et al* (34). GSSH was prepared following the protocol of Luebke *et al* (42). H_2_S_n_ was prepared by a modifying protocol of Xin *et al.* Briefly, 13 mg sulfur powder and 28 mg sodium sulfide were added to 5 ml of anoxic distilled water under argon gas, and the bottle was sealed. The bottle was incubated at 37°C for 24 hours. The pH was adjusted to 9.3 with 6 M HCl, and the solution was diluted to 10 ml with anoxic distilled water. The stock polysulfide concentration was determined with cyanide method and calibrated by using thiosulfate as the standard.

Trifluoromethanesulfonate derivatization and following HPLC analysis was performed following the protocol of a previous report (36).

### Methylene blue method

The methylene blue method was performed as reported previously (32). Two reagents were required, 0.02 M *N,N*-dimethyl-p-phenylenediamine sulfate in 7.2 N hydrochloric acid (DPD reagent), and 0.03 M ferric chloride in 1.2 N hydrochloric acid (ferric chloride agent). Assays are performed in 1.5-mL microfuge tubes. First, 250 μl deionized water was added into a 1.5-ml tube, followed by the addition of 250 μl NaHS, HSSH, or S_8_ solution (in 50 μM, 100 μM, 200 μM, or 500 μM concentration). Second, 20 μl DPD reagent was added, rapidly followed by addition of 20 μl of ferric chloride reagent. The mixture was shaken vigorously and placed in the dark for 30 minutes. After being centrifuged at 17,000 × *g* for 2 minutes. Supernatant (200 μl) was pipetted out and the absorbance at 670 nm was measured. The compounds in supernatant were analyzed using LC-ESI-MS.

### Phylogenetic analysis

The protein sequences of sulfite reductase and cysteine synthase were downloaded from NCBI database (update to 29 August 2024). Redundant sequences were removed using the CD-hit tool with conventional criteria (sequence identity threshold ≥45%, word length =2). The characteristic sequences of the conserved regions were analyzed in the InterPro database. The characteristic sequences of sulfite reductases were PF01077 and PF03460, and the characteristic sequences of cysteine synthases was PF00291. The hmmsearch command in Hmmer tool was used to identify all the featured sequences. Phylogenetic trees were constructed by a neighbor-joining method using ETE3 with a partial deletion, p-distance distribution, and bootstrap at 1,000 repeats. Multiple sequence alignment was performed using MUSCLE.

### Protein expression and purification

Cysteine synthases and sulfite reductases encoding genes were amplified from genomic DNA of *E. coli*, *R. pomeroyi* DSS-3, *S. cerevisiae* S288C, *S. pombe* YHL6381 and *S. islandicus* REY15A genomic DNA. Ferredoxin (Fd, SPO2377), and ferredoxin-NADP reductase (FNR, SPO2637) encoding genes were amplified from genomic DNA of *R. pomeroyi* DSS-3. The expression plasmids (see Table S4) were constructed using primers listed in Table S5. These plasmids were introduced into *E. coli* BL21(DE3) and the strains were incubated in LB medium containing kanamycin (50 μg/ml). When OD_600_ reached 0.6, 0.4 mM isopropyl β-D-1-thiogalactopyranoside (IPTG) was added and the temperature was decreased to 25°C. The cultivation was further continued overnight. Cells were harvested by centrifugation and then re-suspended in binding buffer (200 mM NaCl, 50 mM NaH_2_PO_4_, 20 mM Imidazol, pH 8.0). For protein purification, cell disruption was performed using a Pressure Cell Homogeniser (SPCH-18) at 4°C. Cell lysate was centrifuged to remove the debris and cysteine synthetase/sulfite reductase was purified using the nickel-nitrilotriacetic acid (Ni-NTA) agarose resin (Invitrogen, Waltham, MA, USA). Purification was conducted by following the manufacturer’s instructions. The eluted protein was loaded onto the PD-10 desalting column (GE) for buffer exchange to sodium phosphate buffer (20 mM, pH 7.6). Purity of the proteins was examined via SDS-PAGE.

### Sulfite reductase activity assay in vitro

The assays were performed as previously reported (12). Briefly, for CysIJ, Met5&Met10, and Sir1 activity assays, 0.3 ml reaction buffer (50 mM K_3_PO_4_, pH 7.7, containing 0.1 mM Na_2_EDTA) contained 2 μg/mL purified sulfite reductase, 200 μM sulfite, 100 μM hemin and 2 mM NADPH. The reaction was conducted at 25°C for 10 min. For SPO2634 activity assay, 0.3 ml Tris-HCl buffer (pH 7.5, containing 0.1 M NaCl) contained 2 μg/ml of purified SPO2634, purified ferredoxin (Fd, SPO2377), and purified ferredoxin-NADP reductase (FNR, SPO2637), 200 μM sulfite and 2 mM NADPH. The reaction was conducted at 25°C for 10 min. Produced H_2_S and HSSH were derivatized by mBBr following a previously reported protocol (34), and derivatized products were analyzed by LC-ESI-MS.

As control experiments, 500 μM HSSH was added into 0.3 ml reaction buffer (50 mM K_3_PO_4_, pH 7.7, containing 0.1 mM Na_2_EDTA) with or without 2 mM NADPH. After 10 min incubation at 25°C, produced H_2_S and remaining HSSH were derivatized by mBBr and quantified using RP-HPLC.

### LC-ESI-MS Analysis

The quadrupole time-of-flight, high-resolution mass spectrometer (Ultimate 3000, Burker impact HD, Thermo Fisher, Waltham, MA, USA) was used. The column was InertSustain C18 5 μm (4.6 mm × 250 mm, GL Sciences, Shanghai, China). Injection volume of sample was 10 μl. The source temperature was set at 200°C and the ion spray voltage was at 4.5 kv. Nitrogen was used as the nebulizer and drying gas. The mobile phase was a mixture of pure water and methanol. A linear gradient of solvent A (0.25% acetic acid in ddH_2_O) and solvent B (100% methanol) from 7.5% to 100% to 7.5% (solvent B) in 31 min was used for elution, the column temperature was maintained at 38°C. Data were acquired in full MS (100−1500 m/z) mode. Full mass resolution (40000@ m/z 1222) at full sensitivity.

### Sulfite reductase activity assay in vivo

*E. coli* MG1655-CysJI/Met5&Met10/Sir1 overexpression strains were constructed with pBBR1MCS5 plasmids (Table S4). The strains were cultured in LB medium at 37°C for overnight. The overnight culture (1 ml) was transferred into 100 ml fresh LB medium and cultured to OD_600_=0.4, and then 0.4 mM IPTG was added and the cultivation continued at 37°C for 3.5 h. Cells were collected by centrifugation (4,500×*g*, 10 min) and re-suspended in PBS buffer (pH 7.4, 50 mM) to make a OD_600_=3.0 cell suspension. Sulfite (0.5 mM) was added into 10 ml cell suspension in a 50-ml scale tube. The tube was then incubated at 37□ for 1 h with shaking (200 rpm).

*R. pomeroyi*-SPO2634 overexpression strains were constructed with *R. pomeroyi* DSS-3 Δ*sqr*Δ*fccAB* strain and pBBR1-SPO2634 plasmid (Table S4). Strains were cultured in 1/2 YTSS medium at 30□ for overnight. The overnight culture (1 ml) was transferred into 100 ml of fresh medium and cultured to OD_600_= 0.7, and then 0.1 mM sulfite was added and the cultivation continued until OD_600_ reached 2.0. Cells were collected by centrifugation (4,500×*g*, 10 min) and re-suspended in Tris-HCl buffer (pH 8.4, 50 mM, 20 mM MgCl_2_, 20g/L NaCl) to make a OD_600_=5.0 cell suspension. Sulfite (0.5 mM) was added into 10 ml cell suspension in a 50-ml scale tube. The tube was then incubated at 30□ for 1 hwith shaking (200 rpm).

*S. cerevisiae* S288C-MET5&MET10 overexpression strain was constructed with pJFE3 vector (Table S1). The strain was cultured in SD-URA medium at 30□ for overnight. The overnight culture (1 ml) was transferred into 100 ml of fresh medium and cultured to OD_600_= 0.6, and then 0.1 mM sulfite was added and the cultivation continued until OD_600_ reached 1.0. Cells were collected by centrifugation (4,500×*g*, 10 min) and re-suspended in PBS buffer (pH 7.4, 50 mM) to make a OD_600_=10.0 cell suspension. Sulfite (0.5 mM) was added into 10 ml cell suspension in a 50-ml scale tube. The tube was then incubated at 30□ for 1 h with shaking (200 rpm).

After the incubation, cell suspension was centrifugated (4,500×*g*, 10 min) and 100 μl supernatant was taken after centrifugation. H_2_S, total S^0^, and sulfite concentrations in the supernatant were quantified with following methods.

### H_2_S and sulfite quantification

For quantification of H_2_S and remaining sulfite in obtained supernatant and in aSiRs reaction system, the supernatant and reaction mixture were passed through a 3K Amicon® Ultra centrifugation filter (Sigma-Aldrich, China). The passed-through solution was collected and subjected to mBBr derivatization following protocols reported previously (43–44). Briefly, 30 μl of passed-through solution was added to a PCR tube containing 70 μl of 100mM Tris-HCl buffer (pH 9.5, 0.1mM DTPA), followed by addition of 50 μl of 10 mM mBBr (dissolved in deoxygenated acetonitrile). The reaction was stopped by adding 50 μl of 200 mM 5-sulfosalicylic acid after a 30-minute incubation. All derivatized samples are removed from the hypoxic chamber and stored at 4°C until analyzed by RP-HPLC. Pre-made standard curves of H_2_S and sulfite were used for calculation.

### Total S^0^ quantification

Total S^0^ quantification was performed with the cyanide method (45). First, 0.55 mL 1% boric acid was added into a 1.5-mL tube, heated with boiling water for 1 min and then cooled to room temperature, followed by the addition of 0.25 mL sample (the same volume of deionized water was added to the blank control group). Second, 0.2 mL of 0.1 M potassium cyanide was added, heated with boiling water for 1 min, and then cooled to room temperature. Finally, 0.1 mL Fe(NO_3_)_3_ solution was added and centrifuged at 13000 rpm for 2 min. The supernatant was taken and the absorbance at 460 nm was measured. According to the pre-made standard curve, the detection result is converted to the total S^0^ concentration.

### RP-HPLC Analysis

A C18 reverse phase HPLC column (VP-ODS, 150 3 4 mm, Shimadzu) was pre-equilibrated with 80% Solvent A (10% methanol and 0.25% acetic acid) and 20% Solvent B (90% methanol and 0.25% acetic acid). The column was eluted with the following gradients of Solvent B: 20% from 0 to 10 min; 20%–40% from 10 to 25 min; 40%–90% from 25 to 30 min; 90%–100% from 30 to 32 min; 100% from 32 to 35 min; 100 to 20% from 35 to 37 min; and 20% from 37 to 40 min. The flow rate was 0.75 ml min21. And the bimane adducts were detected by a fluorescence detector (RF-10AXL, Shimadzu) with excitation at 340 nm and emission at 450 nm.

### Cysteine synthase activity assay

Cysteine synthases were expressed in *E. coli* BL21(DE3) (Table S4) and purified with Ni-NTA agarose resin. Activity assay was carried out as described previously. Briefly, 0.3 ml reaction buffer (50 mM PBS buffer, pH 7.4) contained 1 μg/ml purified enzyme, 100 μM pyridoxal phosphate, 1.0 mM–2.0 mM O-acetyl-L-serine (OAS), 1.0–5.0 mM NaSH, GSSH, or Na_2_S_2_O_3_. The reaction was conducted at 30°C for 5 min and the products were derivatized with mBBr and analyzed with LC-ESI-MS. For kinetics assay, the same reaction system contained varied concentrations of NaSH, GSSH or Na_2_S_2_O_3_ (see maintain text). The reaction was conducted at 30°C for 5 min. For the reaction using NaSH as a substrate, the produced cysteine was labeled with mBBr and quantified by HPLC. For the reaction using GSSH or thiosulfate as a substrate, after 5 min reaction, the product was mixed with 0.5 mM TCEP, and the released cysteine was labeled with mBBr and quantified by HPLC. All kinetic parameters were determined using the Michaelis-Menten equation (non-linear regression) embedded in Graphpad prism software.

## Supporting information

including Figure S1-S14, Table S1-S5

## Acknowledgements

This work was supported by the National Key R&D Program of China (2022YFC3401301) and the National Natural Science Foundation of China (92351302).

