## Supplementary material for "Zero valent sulfur is the product of assimilatory sulfite reductase and a substrate of cysteine synthase": including Figure S1-S14, Table S1-S5

Contents:

Figure S1-S14

Table S1-S5

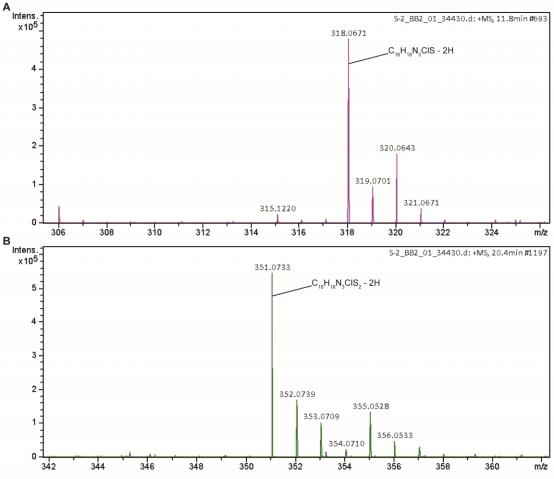

**Figure S1.** MS spectra of **a** (A) and **b** (B) chemicals produced from H2S and Fe^3+^/DMPD.

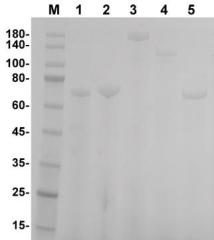

**Figure S2.** SDS-PAGE analysis of purified sulfite reductases/subunits. Lane M, protein

marker (kDa); lane 1, CysI from*E. coli* MG1655; lane 2, CysJ from*E. coli* MG1655; lane 3, MET5 from *S. cerevisiae* S288C; lane 4, MET10 from *S. cerevisiae* S288C; lane 5, SPO2634 from *R. pomeroyi* DSS-3.

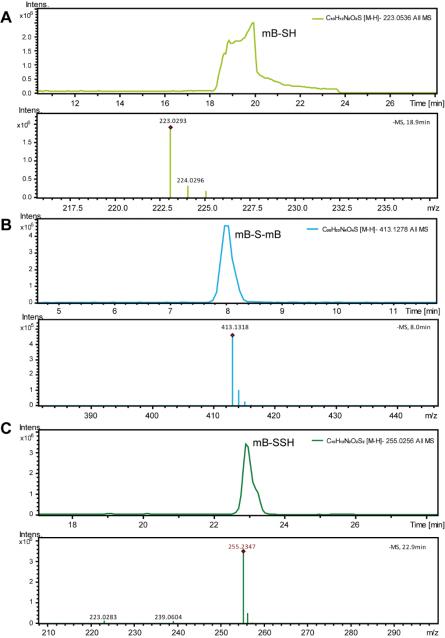

**Figure S3.** LC-ESI-MS analysis of the products from CysIJ catalyzed sulfite reduction. A and B, H2S derivatized by mBBr. C, HSSH derivatized by mBBr.

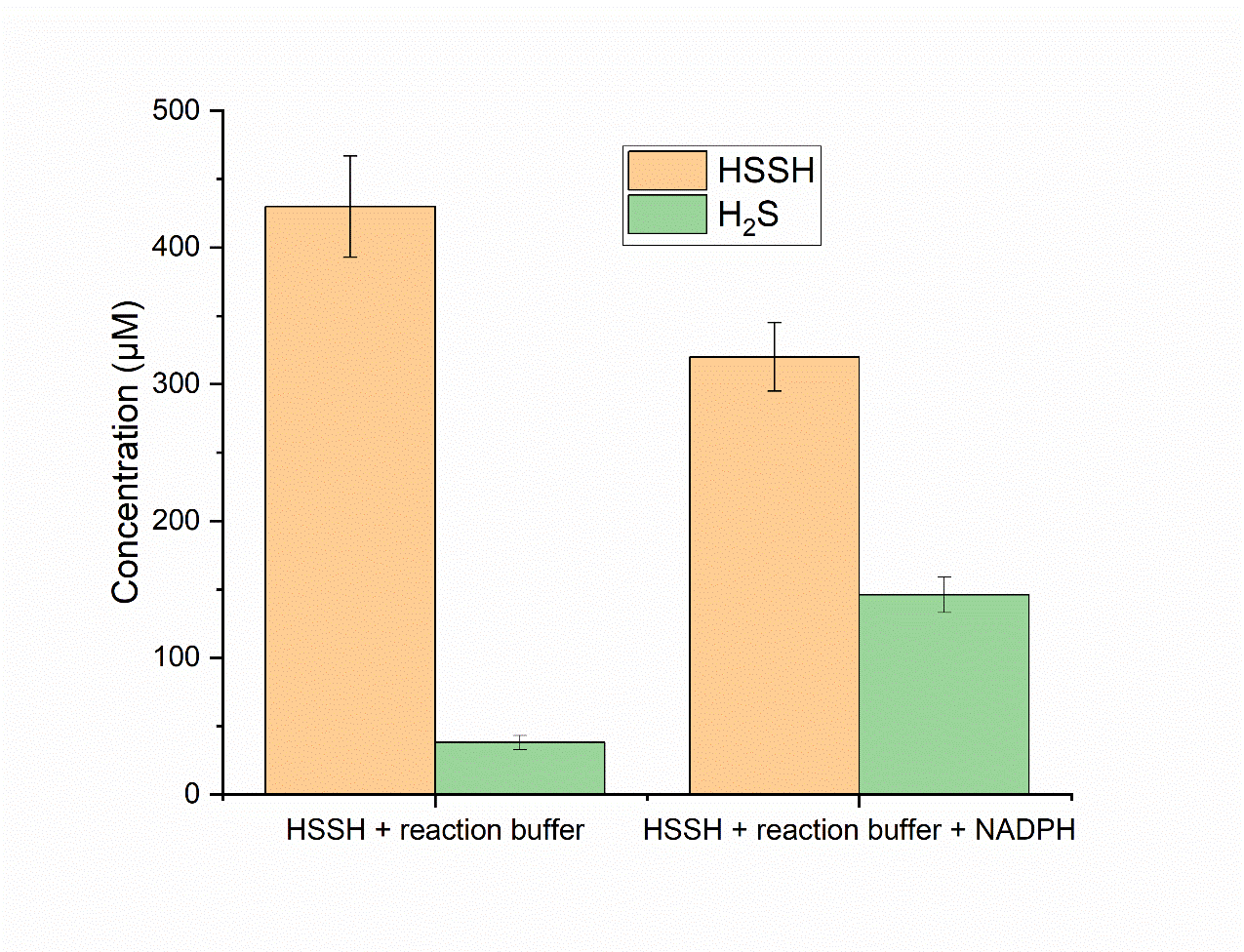

**Figure S4.** HSSH (500 μM) was added into 0.3 ml reaction buffer (50 mM K_3_PO_4_, pH 7.7, containing 0.1 mM Na_2_EDTA) with or without 2 mM NADPH. After 10 min incubation at 25°C, produced H_2_S and remaining HSSH were derivatized by mBBr and quantified using RP-HPLC.

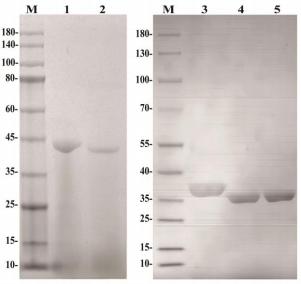

**Figure S5.** SDS-PAGE analysis of purified cysteine synthases. Left panel: lane M, protein marker (kDa); lane 1, CysK from *R. pomeroyi* DSS-3; lane 2, CysM from *R. pomeroyi* DSS- 3. Right panel: lane M, protein marker (kDa); lane 3, CysK from*E. coli* MG1655; lane 4, CysM from*E. coli* MG1655; lane 5, SiRe_1641 from *Sulfolobus islandicus* REY15A.

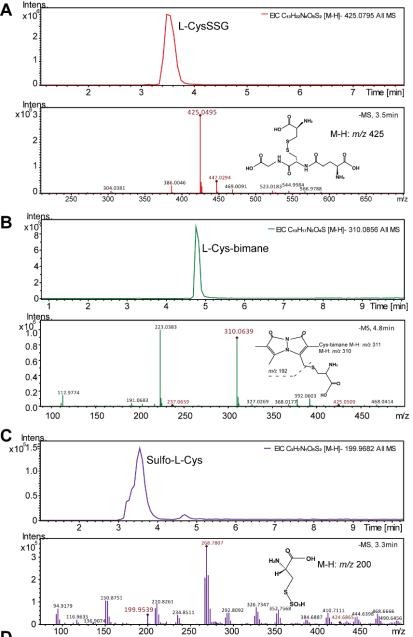

**Figure S6.** LC-ESI-MS analysis of the products from RpCysM-catalyzed reactions. (A) GS- cysteine intermediate was detected when using GSSH as the substrate. (B) L-cysteine was detected when using H2S as the substrate. L-Cys-bimanewas L-cysteine derivatized by mBBr. (C) Sulfo-cysteine intermediate was detected when using thiosulfate as the substrate.

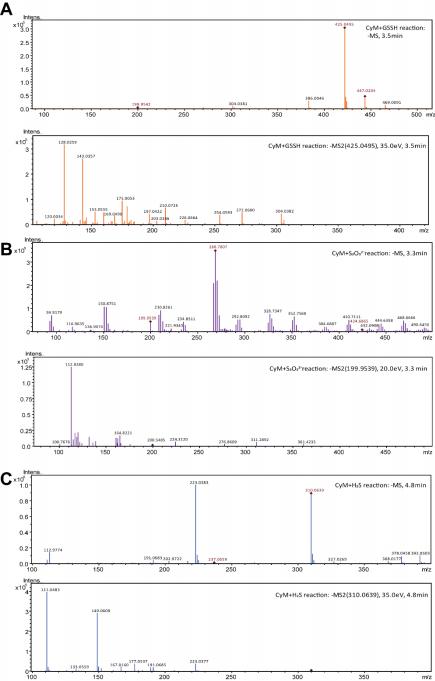

**Figure S7.** LC-MS/MS analysis of the GS-cysteine (A), sulfo-cysteine (B), and mBBr derivatized L-cysteine (C) produced by RpCysM with GSSH, thiosulfate, and H2S, respectively.

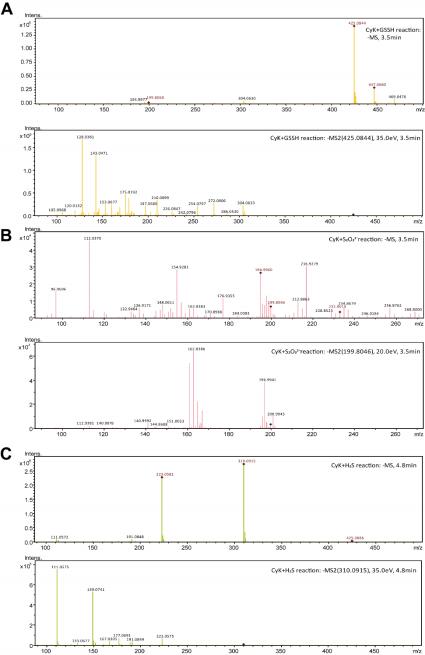

**Figure S8.** LC-MS/MS analysis of the GS-cysteine (A), sulfo-cysteine (B), and mBBr- derivatized L-cysteine (C) produced by RpCysK with GSSH, thiosulfate, and H2S, respectively.

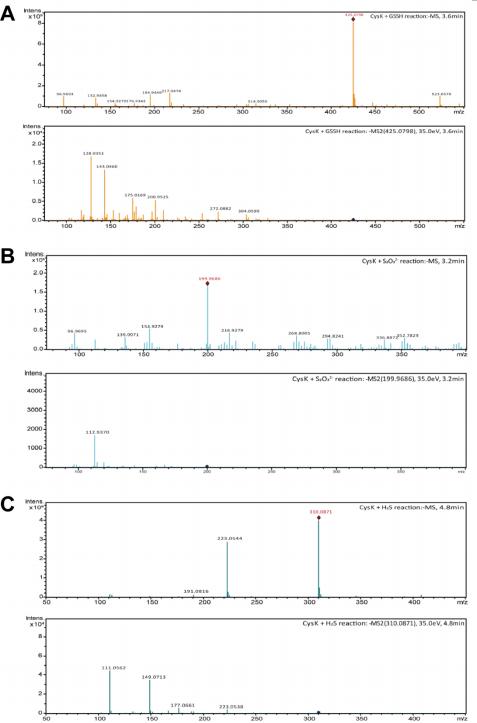

**Figure S9.** LC-MS/MS analysis of the GS-cysteine (A), sulfo-cysteine (B), and mBBr- derivatized L-cysteine (C) produced by EcCysK with GSSH, thiosulfate, and H2S, respectively.

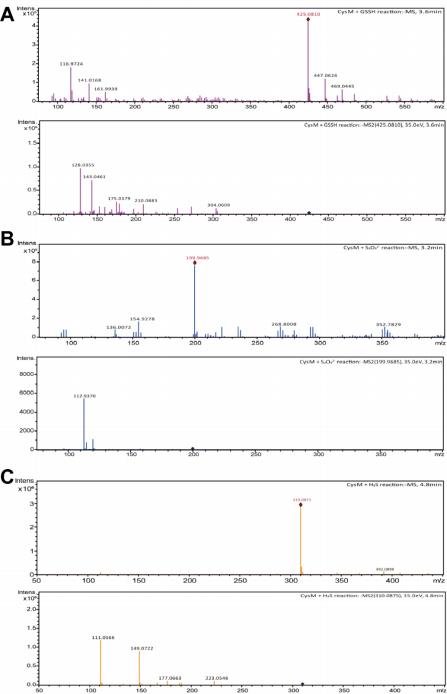

**Figure S10.** LC-MS/MS analysis of the GS-cysteine (A), sulfo-cysteine (B), and mBBr- derivatized L-cysteine (C) produced by EcCysM with GSSH, thiosulfate, and H2S, respectively.

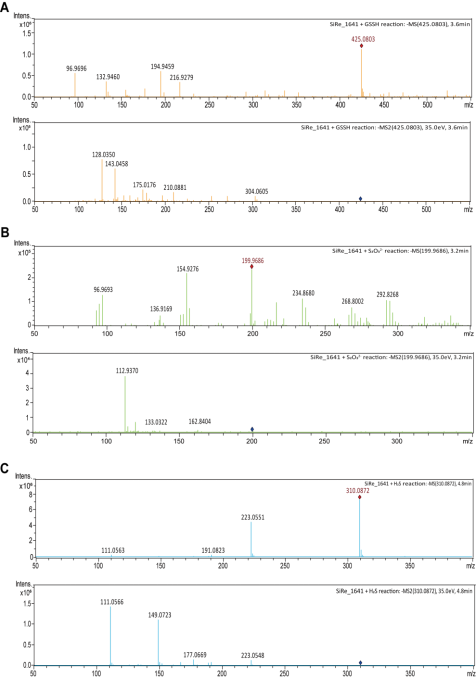

**Figure S11.** LC-MS/MS analysis of the GS-cysteine (A), sulfo-cysteine (B), and mBBr- derivatized L-cysteine (C) produced by SiRe_ 1641 with GSSH, thiosulfate, and H2S,

respectively.

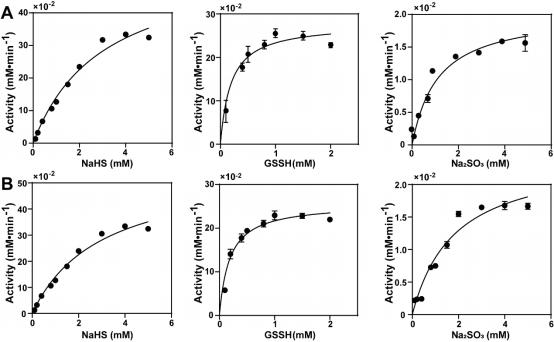

**Figure S12.** Kinetic assays of RpCysK (A) and RpCysM (B) with H2S, GSSH and thiosulfate as substrates. Data were from three independent experiments and presented as average±S.D.

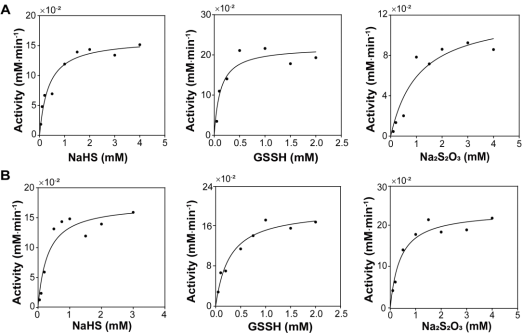

**Figure S13.** Kinetic assays of EcCysK (A) and EcCysM (B) with H2S, GSSH and thiosulfate as substrates. Data were from three independent experiments and presented as average±S.D.

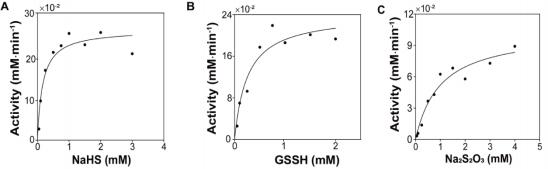

**Figure S14.** Kinetic assays of SiRe_ 1641 with H2S (A), GSSH (B) and thiosulfate (C) as substrates. Data were from three independent experiments and presented as average±S.D.

Table S1. Phylogenetic analysis of 289 sulfite reductases.

| Cluster | ID Node | Organism |
| --- | --- | --- |
| Cluster 1 | WP_014126353.1 KZX14274.1  KAF5420714.1  PPA79744.1  ANF35224.1  EGD18339.1  ABS02816.1  KAJ5553380.1  XP_066694905.1  sp\|P38681.2\|NIR_NEUCR  TPX59622.1 ACZ20059.1 ACQ93374.1 AEX52090.1 TWH58218.1 | Thermoproteus  Methanobrevibacter ANME-2  ANME-2 Bacillus  Xanthomonas Kineococcus  Penicillium  Apiospora  uncultured  Powellomyces Sanguibacter Tolumonas  Rahnella  Desulfitobacterium |
| Cluster 2 | ADN50822.1  WP_221288472.1 AIM27038.1 | Vulcanisaeta  Stygiolobus  Metallosphaera |
| Cluster 3 | QEG41581.1  VAX16676.1  WP_ 188848470.1 WP_ 184679026.1 RIK81626.1  WP_ 197442544.1 TDJ05215.1  TDI80686.1  KAJ9465742.1  KAA6427826.1  KAE8811589.1  KAE8808465.1  KAG6588192.1  GER46021.1  KAK4407409.1  EGB10264.1  XP_005762527.1 KAG7340174.1  KEF43453.1  PNH02096.1  ABG52631.1  WP_299043896.1 WP_097651066.1 | Roseimaritima  hydrothermal  Sulfodiicoccus  Algisphaera  Planctomycetota Lignipirellula  Deltaproteobacteria Betaproteobacteria  Diplonema Trebouxia Hordeum  Hordeum Cucurbita Striga  Sesamum  Aureococcus Emiliania  Nitzschia  Cyanobium Tetrabaena  Trichodesmium uncultured  Candidatus |

|  | WP_291909213.1 WP_276981878.1 GEP46288.1  PSR22557.1  RMH23315.1  QSO52170.1  GIS63460.1  WP_ 171188118.1 WP_ 145358531.1 WP_ 102740259.1 WP_ 197495940.1 | Candidatus  Ferrimicrobium Brevifollis  Sulfobacillus  Acidobacteriota  Alicyclobacillus  Planctomycetaceae  Alienimonas  Alienimonas  Akkermansia  Acidihalobacter |
| --- | --- | --- |
|  | AAC75805.1 | Escherichia |
|  | PRD20934.1  GAK85872.1  GBG34565.1  OLP88546.1  KAF8199148.1  XP_043177304.1 KAI9440511.1  OCB89998.1  KAK7061135.1 KAI3608505.1 | Trichonephila Vibrio  Hondaea  Symbiodinium Pholiota  Rhizoctonia  Lactarius  Sanghuangporus Paramarasmius Moniliophthora |
|  | CAB11176.1 | Schizosaccharomyces |
|  | VUG16533.1 | Brettanomyces |
|  | NP_ 116579.1 | Saccharomyces |
|  | KAG0678211.1 GCE97350.1  SLM39464.1  XP_049151562.1 KAF3401165.1  KAA0178054.1 POM69529.1  XP_004990079.1 KAI8807475.1  KAI9353484.1 OBZ88583.1  KAJ1979091.1 OMH81431.1 KAJ2009528.1 | Kluyveromyces  Zygosaccharomyces Lasallia  Colletotrichum Talaromyces  Cafeteria  Phytophthora Salpingoeca  Cladochytrium Obelidium  Choanephora Dimargaris  Zancudomyces Coemansia |
| Cluster 4 | WP_091792141.1 ODA68264.1  GAA5654175.1 EJM99334.1 | Peptococcus  Methyloligella Brucella  Phyllobacterium |

QFT44903.1

ASL48163.1

KEZ78409.1

AFY74132.1

EKQ68488.1

AFZ04234.1

AFY47725.1

QDU33230.1

WP_292655332.1 WP_353663014.1 WP_299998028.1 WP_305532260.1 ADC89957.1

WP_297525046.1 WP_201335301.1 WP_200673635.1 WP_201352181.1 WP_ 129104492.1 WP_ 172128353.1 APZ96131.1

WP_290623662.1 KYH39988.1

KKK43102.1

WP_014449279.1 WP_ 151619990.1 WP_018701792.1 GAA5528742.1

WP_205839031.1 WP_205007292.1 WP_057003017.1 AEV69918.1

EHJ00794.1

SMC05887.1

GAA5528643.1

WP_ 173730930.1 AFZ14489.1

WP_079255002.1 ELY78545.1

WP_202953832.1 ADB61906.1

WEL16481.1

ADO44842.1

WP_259099578.1

Pseudomonas

Burkholderia

Salinisphaera

Synechococcus

Leptolyngbyaceae Calothrix

Nostoc

Poriferisphaera Nitratifractor

Hydrogenimonas Sulfuricurvum

Sulfuricurvum

Thermocrinis

Sulfurovum

unclassified

Persephonella

Hydrogenimonas Arcobacter

Arcobacter

Fuerstiella

unclassified

Candidatus

Candidatus

Leptospirillum Heliorestis

Anaeromusa

Herpetosiphon Bilifractor

Sporolactobacillus Agrilactobacillus	Acetivibrio

Clostridium

Sulfobacillus

Herpetosiphon

Calidifontibacillus

Crinalium

Halococcus

Natrinema

Solirubrobacter Haloterrigena	Halorhabdus

Hydrogenobacter Candidatus

|  | ABL83293.1  WP_320051281.1 WP_289401893.1 WP_235826971.1 WP_ 129103884.1 WP_229234160.1 QDU70873.1  KAF0248333.1  BCM90664.1  WP_299370467.1 GAA3767814.1  WP_ 143720194.1 WP_304045879.1 SIL51448.1  WP_303946192.1 WP_ 165134532.1 WP_036302080.1 WP_326926227.1 VGO19700.1  WP_ 170291659.1 CAG0992393.1  WP_040199035.1 WP_282002389.1 QND78565.1  ATY93604.1  AHL31010.1  AAF27543.1  BAU56604.1  WP_200195964.1 WP_292832310.1 SDD95998.1  WP_ 134286370.1 WP_ 114899022.1 WP_371295956.1 GEN04639.1  SDD96082.1 | Nocardioides uncultured  Sulfurovum  Aliarcobacter Arcobacter  Candidatus  Mucisphaera  bacterium  Abditibacteriota Edaphobacter  Terriglobus  Deinococcus  Jatrophihabitans Mycobacteroides Corynebacterium Microbacterium Microbacterium Candidatus  Pontiella  Heliobacterium  Planctomycetaceae Geoalkalibacter  Geotalea  Candidatus  Candidatus  Candidatus  Allochromatium Halorhodospira Halorhodospira Mesorhizobium Cupriavidus  Pseudomonas Aquirhabdus Burkholderia Acetobacter  Cupriavidus |
| --- | --- | --- |
|  | AAV95879.1 | Ruegeria |
|  | WP_294105533.1 WP_325590877.1 WP_303054586.1 OQC31671.1  ADO82703.1  WP_297406306.1 | Sphingomonas Iamia  Victivallis  Verrucomicrobia Ilyobacter  uncultured |

WP_332478929.1 OQA01240.1

OQB78970.1

WP_291281609.1 WP_289221079.1 KXB85311.1

WP_020310923.1 KXB88425.1

WP_274941657.1 WP_337766396.1 WP_349011068.1 CVI64900.1

CUN04777.1

SCI34541.1

WP_262070057.1 WP_013655115.1 WP_317731059.1 WP_307993834.1 AGK98990.1

PRR83705.1

WP_ 162369330.1 KXB64529.1

EJU07917.1

ACZ07360.1

OHW61533.1

WP_048569142.1 WP_019878989.1

SCN26124.1

UMZ72693.1

WP_265823795.1 CAK8715638.1

CAK8719475.1 TKJ34400.1

TDX50979.1 SJZ78723.1	AEJ61941.1

WP_315015124.1 TLD95584.1

SDG26476.1 VAX36249.1 BBM89323.1 SEQ15256.1 BBE19865.1

Cetobacterium	Planctomycetes Planctomycetes Fusobacterium	Faecalibaculum Veillonella

Megasphaera Veillonella

Chordicoccus

Phascolarctobacterium Mediterraneibacter

Eubacteriaceae

Anaerobutyricum

uncultured

Ohessyouella

Cellulosilyticum Anaerocolumna

uncultured

Clostridium

Clostridium

Anaerotalea

Leptotrichia

Fusobacterium Sebaldella

Andreesenia

Clostridium

Succinispira

Clostridium

Natranaerofaba Geovibrio

Candidatus

Candidatus

Planctomycetes Orenia

Selenihalanaerobacter

Spirochaeta uncultured

Helicobacter	Bacteroidales hydrothermal Spirochaetota Segatella

Aquipluma

|  | VAX22157.1  GAT35297.1  WP_331016035.1 WP_235969639.1 WP_293270630.1 EME71902.1  WP_332894804.1 KAF0191654.1  WP_325328049.1 RLL51003.1  RMH52625.1  EDM25005.1  SNZ11452.1  AEE53707.1  WP_077130662.1 RAI99451.1  SDM70161.1  GBD25812.1  EYF06509.1  GBD02918.1  CUV02505.1  WP_337863914.1 WP_238443600.1 KLR62045.1  CAI2718116.1  CBK40184.1  CAI8024908.1  WP_254010452.1 WP_206292550.1 OGQ93275.1 | hydrothermal  Terrimicrobium Candidatus  Anaeromyxobacter Nannocystis  Paramagnetospirillum Magnetovibrio  Gammaproteobacteria Sulfuricella  Mariprofundus  Zetaproteobacteria Lentisphaera  Persephonella  Haliscomenobacter  Spirosoma  Chitinophaga Daejeonella  bacterium  Chondromyces bacterium  hydrothermal Nitrososphaera  Salsipaludibacter  Actinobacteria Nitrospina  Nitrospira Geodia  Limnofasciculus Humisphaera  Deltaproteobacteria |
| --- | --- | --- |
| Cluster 5 | RJP23896.1 PIP67686.1  OGX25733.1 RJQ28770.1 VHV38821.1 SUG71549.1 BCO10446.1 AEH44006.1 KNZ68571.1 ACX52333.1 SMC00018.1 AEG16294.1 AGL03340.1 | Candidatus  Candidatus  Omnitrophica Peptococcaceae Clostridioides  Salmonella  Desulfolithobacter Thermodesulfatator Thermincola  Ammonifex  Thermanaeromonas Desulfofundulus  Desulfoscipio |

|  | SNQ61253.1 | Candidatus |
| --- | --- | --- |
|  | BDQ38728.1 PLX47257.1 | Pseudodesulfovibrio Desulfobulbaceae |
|  | WP_ 108308452.1 | Thermodesulfobium |
|  | BDQ36097.1 PWR72355.1 | Pseudodesulfovibrio Methanospirillum |
|  | OIO04147.1 AIF54213.1 | Desulfovibrionaceae Pelosinus |
|  | AGB41179.1 AEA46668.1 OAM92504.1 SUP43984.1 | Halobacteroides Archaeoglobus Pelosinus  Veillonella |
|  | ADC69017.1 EHP84882.1 GBD99048.1 GBE41111.1 | Methanocaldococcus  Methanotorris bacterium  bacterium |
|  | WP_278742369.1 | Megamonas |
|  | GAW91740.1 GBE17736.1  KAF5036219.1 | Calderihabitans archaeon  anaerobic |
|  | AET66482.1 ACL17181.1 ACD22888.1 WLD26378.1 | Desulfosporosinus Methanosphaerula Clostridium  Clostridioides |
|  | GAV22117.1 KJJ84223.1  TLU57669.1 ABB24416.1 GFO97279.1 KNZ70575.1 XER07471.1 XER11215.1 | Carboxydothermus  Candidatus Chlorobium Pelodictyon groundwater Thermincola Sporomusa Sporomusa |

Note: 4 representative enzymes are shown with underline.

Table S2. Phylogenetic analysis of 735 cysteine synthases.

| Cluster | ID Node | Organism |
| --- | --- | --- |
| Cluster 1 | WP_313561741.1 KNY26424.1  CAH2942971.1  PHM43162.1  KKR69583.1  WP_244145184.1 CZF78179.1  WP_317535548.1 GJJ35119.1  WP_024805385.1 WP_356995616.1 WP_040750577.1 GFI23655.1  GFI29825.1  WP_045875124.1 WP_ 165913567.1 TCO88741.1  CCO22305.1  WKZ48475.1  KFC17095.1  WP_274076748.1 WP_344660838.1 PZU93388.1  QEI39871.1  GEN26540.1  WP_328239200.1 WP_ 184694028.1 WP_ 189056211.1 AKT40690.1  CCF12539.1  SCM11940.1  RZO31370.1  WP_025354931.1 WP_218476955.1 WP_ 102387923.1 WP_206126043.1 RLA43414.1  WP_359361219.1 WP_363228853.1 SEG46938.1  WP_277960392.1 XDL19947.1 | Ruminiclostridium Pseudobacteroides  uncultured  Xenorhabdus  Parcubacteria  Paraburkholderia Grimontia  Nitrosomonas  Corynebacterium Nocardia  Nocardia Nocardia  Lachnospiraceae Lachnospiraceae Pseudofrankia  Marichromatium Chthoniobacter  Maridesulfovibrio Anaerolineales  Cutibacterium  Pseudomonas  Catenulispora  Pseudanabaena  Dolichospermum Halovibrio  Shouchella  Saccharothrix  Longimycelium Chondromyces  Brevibacillus Bacillus  SAR116 Kutzneria Nocardia Vibrio  Burkholderia  Gammaproteobacteria  Streptomyces  Streptomyces  Paenibacillus  Staphylococcus Dickeya |

TCU50258.1

EYF05055.1

ERJ97380.1

CUP19573.1

CAC9975516.1 SEE22512.1

AEV98680.1 BAV05925.1

WP_369616568.1

RAJ29101.1 AOE92688.1 PVZ05892.1 PRY88556.1 ORL43814.1 TCO11085.1

TYP69968.1

SEC29777.1

SLM92927.1

SDK99161.1

KPC80740.1

EAQ80544.1

WP_356556387.1 ATL31989.1

PPT12123.1

RSN29095.1

QES47491.1

WP_ 145772533.1 SES35060.1

WP_344163106.1 PZW20782.1

WUX35132.1

WSX26215.1

WP_261585165.1 SIJ91699.1

SUA79628.1 EPJ84757.1

GBH18367.1 EPN23876.1 SDY43573.1 SMR71958.1 SNY97082.1 RMR09649.1 TDV51888.1 OKH92139.1

Curtobacterium Chondromyces Ruminococcus Agathobacter

Flavobacterium Tenacibaculum Niastella

Filimonas

Flavobacterium Pedobacter

Ralstonia

Actinomycetospora

Mongoliibacter Zunongwangia

Natronoflexus

Aquimarina

Arthrobacter

Brachybacterium Paenibacillus

Streptomyces

Blastopirellula

Streptomyces

Streptomyces

Streptomyces

Amycolatopsis

Streptomyces

Micromonospora Lentzea

Nocardiopsis

Thermosporothrix Streptomyces

Streptomyces

Planotetraspora	Mycobacteroides Pandoraea

Pseudomonas

Pseudomonas

Pseudomonas

Micromonospora Aliiroseovarius

Halomonas

Pseudomonas

Actinophytocola Streptomyces

|  | RZU18021.1 PBC86986.1 | Streptomyces Streptomyces |
| --- | --- | --- |
| Cluster 2 | PJE02371.1  WP_ 193825512.1 WP_010419882.1 ALU11385.1  WP_ 125740919.1 ALL00712.1  RLE56066.1  WP_008083777.1 | Leptospira  Leptospira  Leptospira  Ignicoccus  Candidatus  Pyrodictium Thermoprotei Candidatus |
| Cluster 3 | CEG65771.1  XP_055350340.1 TFG35403.1  EEC05264.1 KOB51926.1 PIM94780.1 PIM96424.1 PIM96950.1  WP_237558828.1 SMC78645.1  EEG89274.1  KAI0215617.1 KAJ1671120.1 PRD18789.1  SJM66503.1  KPA13804.1  EDY15779.1  EID77315.1  EPR29837.1  EPD10039.1  XP_009652046.1 GAE76007.1  OQA37428.1  WP_207862935.1 WP_254061617.1 CAA3816734.1  CAA2953972.1  ERH08995.1  WP_224828057.1 WP_315908517.1 SEN27780.1  WP_266013030.1 WP_ 121727239.1 | Rhizopus  Paramacrobiotus Parcubacteria  Ixodes  Operophtera  Candidatus  Candidatus  Candidatus  Candidatus  Oscillospiraceae Coprococcus  Lamellibrachia  Coemansia  Trichonephila  Frigoribacterium Candidatus  Chthoniobacter Rhodococcus  Geobacillus  Lacticaseibacillus  Verticillium  Cutibacterium Acidobacteria  Acanthopleuribacter Granulicella  Staphylococcus  Olea  halophilic  Saliphagus Halapricum Prevotella  unclassified Helicobacter |

PPV18032.1

AGF55535.1

ARQ07885.1

QAS23049.1

OCX49408.1

XP_035449125.2 CDO76302.1

QRW10917.1

GAA53379.1

XP_001390590.3

sp|Q2V0C9.1|CBS_APIME	sp|G5EFH8.1|CBS1_CAEEL

JAA48330.1

WP_326123129.1 WP_252506913.1 WP_021191817.1 TJZ52531.1

QBR11433.1

TFH60907.1

WP_367167674.1 WP_282498408.1 SQI42233.1

WP_230372930.1 WP_ 126803636.1 AEV26414.1

WP_ 178335601.1 WP_338008169.1 WP_277552377.1 ABC45805.1

WP_ 137314544.1 QQR62842.1

ETO92078.1

WP_ 192577603.1 QEN44469.1

WP_216374074.1 WP_ 171597272.1 WP_ 155135183.1 VFK73321.1

WPD21554.1

XP_002502084.1 GBG34329.1

KAK7242038.1 EDM77348.1

Clostridium

Clostridium

Macrococcoides

Lactiplantibacillus Ligilactobacillus	Spodoptera

Trametes

Ceratobasidium Clonorchis

Aspergillus

uncultured

uncultured

Desmodus

Metasolibacillus Anoxybacillus

Sphingobacterium Sphingobacterium Sphingobacterium

Candidatus Erwinia

Pantoea

Serratia

Enterobacter

Aliidiomarina Azospira

Natronomonas

Natronoglomus Halobaculum

Salinibacter

Pseudoduganella

bacterium

Legionella

Francisella

Francisella

Francisella

Marinifilum Roseibium

Candidatus

Candidatus

Micromonas Hondaea

Aureococcus Plesiocystis

|  | AEX85522.1  ALM75563.1  KAA0210709.1 ABL66272.1  RMG11111.1  CUS55351.1  WP_ 143381964.1 WP_020538003.1 WP_304200992.1 EJF52796.1  KXK46614.1  WP_304992618.1 WP_273897731.1 WP_ 198023164.1 BBA95580.1  WP_094723818.1 ADP84692.1  EIC68435.1  TWS19810.1  WP_ 198428424.1 WP_ 157180386.1 WP_003797493.1 WP_ 146321028.1 WP_214154474.1 | Marinitoga  Thermococcus  Chlorobiota  Chlorobium  Planctomycetota hydrothermal  Formosa  Lewinella  Flavobacterium Saprospira  Bacteroidetes  Burkholderia  Pseudomonas  Mesorhizobium Streptomyces  Rhodococcus Pseudofrankia  Mycobacteroides  Tsukamurella Nocardia  Protofrankia Arthrobacter Humibacter Kineosporia |
| --- | --- | --- |
| Cluster 4 | RNC81489.1 UPT75414.1 RLI10059.1  TEU08362.1  GHT01053.1  GBF27531.1  WP_058983531.1 WP_089788918.1 XP_005772444.1 AWE41469.1  RHV79075.1  WP_242879328.1 QCV87897.1  KYC82915.1  SLM91065.1  WP_273565103.1 GAU86451.1  PXV81572.1 PPR16882.1 | Phycisphaera  Elusimicrobiota  Candidatus  Candidatus  Synergistales  bacterium  Halobacterium Halobiforma  Emiliania  Actinobaculum Clostridium  Butyrivibrio  Acidipropionibacterium  Heyndrickxia  Brachybacterium Methylobacterium Bosea  Nitrosomonas  Pseudomonadota |

|  | SDJ86680.1 AFU01937.1 ADI05365.1 TCT16092.1 PJA73338.1 PKQ15274.1 TVQ76568.1 GBE21923.1 RMG93086.1 KTB49298.1 BAB59731.1 CAC11675.1 EQD35550.1 EQB67198.1 UCC57706.1 SPC33587.1  WP_337863426.1 TLX94936.1  WP_291765575.1 PVU73710.1  GGT93808.1 | Nocardioides Nocardia  Streptomyces  Natranaerovirga Nitrospirae  Actinobacteria  Phycisphaeraceae bacterium  Candidatus  Dehalogenimonas Thermoplasma  Thermoplasma mine  Thermoplasmatales  Candidatus  Candidatus  Nitrososphaera Nitrososphaerota Caldivirga  Sulfolobales  Sulfodiicoccus |
| --- | --- | --- |
|  | ADX85705.1 | Sulfolobus |
|  | BDR93349.1  ABN69785.1  PNV80199.1  AET33849.1  BBE42620.1  ESQ21196.1  WP_292320782.1 BBK26391.1  BBE42199.1  ALL00818.1  AFL66531.1  UCG41127.1  CQD24773.1  WP_296388246.1 KAA1280543.1  RLT55758.1  TMK26254.1  TFH17522.1  TML25775.1  WP_236601796.1 KRT69075.1 | Vulcanisaeta  Staphylothermus Fervidicoccus  Pyrobaculum  Conexivisphaera uncultured  Caldisphaera synthetic  Conexivisphaera Pyrodictium  Desulfurococcus Acidimicrobiia  Mycolicibacterium Williamsia  SAR202  Chloroflexota  Actinomycetota Acidimicrobiales Actinomycetota Ktedonobacter  candidate |

|  | BBM89228.1 EGQ61061.1 GIS46408.1 GBF89740.1 OLQ08845.1 STW05514.1 SUG71403.1 | Spirochaetota  Acidithiobacillus  Gammaproteobacteria  Raphidocelis Symbiodinium Klebsiella  Salmonella |
| --- | --- | --- |
|  | AAC75474.1 | Escherichia |
|  | PWI48476.1  GBL17251.1  TMG39664.1  GIW70681.1  PYS32010.1  CEK14978.1  WEK37920.1  OJX37687.1  OQB17526.1  WP_080803637.1 WP_092229638.1 WP_258930926.1 AEH22379.1  WP_045218853.1 WP_092062550.1 WP_325644966.1 WP_ 171267194.1 BES64211.1  WP_ 191055911.1 WP_ 136989937.1 WP_256045495.1 RPG05710.1  RUM59827.1  WP_324715852.1 KUG04260.1  PYN06185.1  UCH37864.1  PHX72826.1  PYT12741.1  WP_ 132286832.1 OYV05788.1  SFH99120.1 QJW95035.1 BDC48637.1 CCH48467.1 | Candidatus  Chloroflexota  Chloroflexota  Planctomycetota Acidobacteriota Chthonomonas Candidatus  Flavobacteriia  Deltaproteobacteria Desulfamplus  Desulfobacula  Desulfoferrobacter  Thermodesulfobacterium  Desulfonatronum  Desulfonauticus  Humidesulfovibrio Oceanidesulfovibrio Gottschalkiaceae  Planktothrix  Martelella  Stutzerimonas  Pelagibacteraceae Persephonella  Limnochorda hydrocarbon Candidatus  Candidatus Opitutia  Acidobacteriota Kribbella  Verrucomicrobiales  Planctomicrobium Frigoriglobus  Bryobacterales  Pseudodesulfovibrio |

|  | TRO53691.1  WP_300061268.1 WP_243638211.1 TRZ51159.1  GIR28323.1  UCD96303.1  RMF06434.1  UCD02627.1  CCY04062.1  KXU52325.1  WP_312032262.1 WP_323723391.1 CDD91530.1  AAL95416.1  WP_089758297.1 KAH0576451.1  CAL6019137.1 VFK16175.1  PIZ72466.1  RZD15511.1  RLF73289.1  ABN69466.1  CDD22689.1  AZV47254.1  WP_325276649.1 PWM77138.1  CDC70929.1  UTW42251.1  WP_233998229.1 OGT94700.1  WP_ 154178644.1 PRD31979.1  AGK96672.1  BDC92876.1  WP_346687505.1 WP_341418758.1 WZS84515.1  WP_089528461.1 WP_371262088.1 ESX60542.1  CAK9100958.1 | Candidatus  Thermomonas  Rubrobacter  Dehalococcoidia Euryarchaeota  Candidatus  Candidatus Candidatus  Faecalibacterium  Candidatus  Hujiaoplasma Veillonella  Coprobacillus Fusobacterium  Halarsenatibacter  Spironucleus Hexamita  Candidatus  Candidatus  Candidatus  Thermococci  Staphylothermus  Firmicutes Nautilia  Romboutsia  Clostridium  Staphylococcus bacterium  Erythrobacter  Gemmatimonadetes  Larkinella  Trichonephila Clostridium  Treponema Enteroscipio Paenibacillus Vibrio  Pantoea  Rhizobacter  Mesorhizobium Durusdinium |
| --- | --- | --- |
|  | AAV95512.1 | Ruegeria |
|  | ORY28396.1 | Rhizoclosmatium |

OLP84007.1 OLP98525.1 OLP79393.1

OLQ02765.1

VEB44754.1

RMH57771.1

WP_082614592.1 WP_076768479.1 RAV78582.1

PKZ21419.1

PMC80688.1

OYQ68070.1

WP_260853922.1 WP_317263764.1 PIO61026.1

UKI20373.1

WP_054494025.1 TAK83174.1

UCC93303.1

VTT88391.1

ELZ00631.1

PSQ22461.1

WP_370240205.1 ABG40402.1

WP_242982837.1 WP_334390599.1 PKP02734.1

QCC85816.1

PWL64378.1

QDU63159.1

WP_318623234.1 WP_356218922.1 KPI23437.1

OEU07594.1

WP_ 192533053.1 TMD56937.1

KAJ4702416.1

GAY54168.1

RWR79411.1

KAK1286664.1

EMS49642.1

XP_040248276.1 URD87860.1

Symbiodinium

Symbiodinium

Symbiodinium

Symbiodinium

Chromobacterium Candidatus

Ligilactobacillus Jeotgalibaca

Aerococcus

Aerococcus

Aerococcus

Aerococcus

Paraburkholderia Rhodovulum

Teladorsagia

Oscillospiraceae Ardenticatena

Betaproteobacteria Thermoplasmata

Halorubrum

Natrialba

Halobacteriales

Marisediminitalea Paraglaciecola

Faecalibacterium Bradyrhizobium Bacteroidetes

Desulfovibrio

Desulfovibrionaceae Planctomycetes

Mycolicibacterium

Streptomyces	Actinobacteria Fragilariopsis	Phaeovibrio

Chloroflexota Melia

Citrus

Cinnamomum

Acorus

Triticum Aegilops Musa

|  | KAF3333414.1  PWZ05116.1  OEL19139.1  WP_346375223.1 OEU73014.1  VPC27650.1  CGG84900.1  KAH3732784.1 KAH1202607.1 KAF8732891.1 QHO19637.1  OEL30495.1  AGT02607.1  KAJ9440351.1 KIH61266.1  KAI6179209.1 PIO71181.1  XP_001733225.1 WP_320596381.1 KAF0229095.1  GJG85719.1  WP_305923677.1 PLT87888.1  TXT27880.1 | Carex  Zea  Dichanthelium Aetherobacter  Desulfuromonadales  Streptococcus Streptococcus Pelomyxa  Glycine Digitaria Arachis  Dichanthelium Strigomonas  Diplonema  Ancylostoma  Aphelenchoides Teladorsagia  Entamoeba  Absicoccus  Fusobacteriota  Gemmatimonadetes  Nonomuraea  Sinorhizobium  Planctomycetota |
| --- | --- | --- |
|  | AAC75467.1 | Escherichia |
|  | CDH70688.1  KAG7372636.1  WP_082704188.1 KAH1237418.1  XP_030943452.1 XP_052193824.1  KAE8769916.1 CCA17822.1  TFJ81847.1  OUW47848.1  BFI31968.1  KAF8066211.1 PRW44930.1  KAF8067153.1 KAG7674702.1 PRW58032.1  PHT41250.1  KAH9754334.1 | Pseudomonas Nitzschia  Paenibacillus Glycine  Quercus Diospyros Hordeum Albugo  Nannochloropsis Synechococcus  Marchantia  Scenedesmus Chlorella  Scenedesmus Chlorella  Chlorella Capsicum Citrus |

|  | KAF4374418.1  PWA90620.1  GAY54164.1  KHN42175.1  KAJ6855009.1  XP_024957103.1 KVH91024.1  EMS61355.1  KAF8655790.1 KAE8733417.1 QCD91844.1  KAG6586292.1 KAL0368100.1 | Cannabis Artemisia Citrus  Glycine Populus Citrus  Cynara  Triticum Digitaria Hibiscus Vigna  Cucurbita Sesamum |
| --- | --- | --- |
| Cluster 5 | WP_296411095.1 WP_051160992.1 WP_086864998.1 WP_ 163371444.1 SFU29001.1  VVE58120.1  RKD84646.1  SDV50747.1  WP_360796063.1 WJN59185.1  WP_281648632.1 GHT78541.1  WP_ 109743779.1 WP_035219688.1 GED15767.1  WP_285012860.1 KZC01280.1  KIG14946.1  CAB9509284.1  WP_263781152.1 WP_364114148.1 PYQ68214.1  XP_001396038.3 EER10573.1  WP_ 127022753.1 TML73599.1  RJQ82700.1  PYO76422.1  KAK5988894.1 GAA1261043.1 | Zoogloea  Nocardia  Amycolatopsis  Endozoicomonas Aliiroseovarius  Pandoraea  Kushneria  Chitinasiproducens Actinosynnema  Pseudomonas  Parendozoicomonas Bacteroidia  Arcicella  Desulfatibacillum Aneurinibacillus Oligella  Methylobacterium  Enhygromyxa Seminavis  unclassified  Actinoplanes  Acidobacteriota Aspergillus  Perkinsus  Flagellimonas Actinomycetota Amycolatopsis  Gemmatimonadota Cladobotryum  Oryzihumus |

|  | UCC84297.1  PYP86874.1  WP_208451094.1 WP_089383865.1 WP_ 139176564.1  PIB01911.1  KKY14223.1  KAK9857765.1 KAF4494306.1  KAG7150165.1 XP_004333552.1 XP_004998218.1 TFJ84572.1  KAJ9466184.1  KAH7501026.1 CCA16821.1  OQS04529.1  KOO23676.1  XP_004354329.1 KAA0160754.1 KAA0149527.1 KAI9011644.1  WP_ 101039182.1 WP_ 185817675.1 ONH66201.1  XP_026191563.1 CEL71048.1  KAF0119325.1 KAG1655709.1 GAD55453.1 | Gemmatimonadota  Blastocatellia Burkholderia Halorubrum Jannaschia  Cercospora Diplodia  Penicillium Fusarium  Verticillium  Acanthamoeba  Salpingoeca  Nannochloropsis Diplonema  Phytophthora Albugo  Thraustotheca  Chrysochromulina Cavenderia  Cafeteria Cafeteria  Hyaloraphidium Psychromonas Shewanella  Cyberlindnera Cyclospora  Neospora  Xanthobacteraceae  Nymphon  Limimaricola |
| --- | --- | --- |
|  | AAV95198.1 | Ruegeria |
|  | TLY85835.1  KAK2560495.1 ONH73330.1  KAJ3372763.1 | Gammaproteobacteria Acropora  Pichia  Kappamyces |
|  | CAA19052.1 | Schizosaccharomyces |
|  | KAI3619846.1  OCB86888.1  SPO26108.1  XP_018001747.1 KAF5244390.1  KPM41549.1  KAJ6446508.1 | Moniliophthora Sanghuangporus  Ustilago  Phialophora Fusarium  Neonectria  Purpureocillium |

|  | KND94675.1  XP_043181979.1 KAF0852140.1  KAH8067748.1 CAK9013900.1  OLQ12318.1  KAK1743413.1 GAX26691.1  KAG7358090.1 XP_017995599.1 KAI5906473.1  DBA02402.1  KAK1939373.1 KAA0165429.1 XP_007512946.1 KAF8063062.1  QKY15310.1  XP_024520240.1 KAF3780173.1  XP_006474900.1 XP_031096819.1 KAK4439705.1 RKP22983.1  KAJ2013480.1  KAJ2086887.1  XP_014170333.1 XP_016583295.1 KJZ77448.1  KAI1251649.1  KAJ9625740.1  RMZ66893.1  XP_025566059.1 EGU12017.1  KAI5480163.1  XP_052942254.1 KAG2016210.1 KAJ8695927.1  XP_060122916.1 XP_056062159.1 CEH12592.1  KAK0522778.1 KAK0565969.1 | Tolypocladium Rhizoctonia  Andalucia  Aureococcus  Durusdinium  Symbiodinium Skeletonema  Fistulifera  Nitzschia  Phialophora Candida  Lagenidium Phytophthora Cafeteria  Bathycoccus Scenedesmus Polytomella Selaginella  Nymphaea  Citrus  Ipomoea  Sesamum  Syncephalis Coemansia  Coemansia  Grosmannia Sporothrix  Hirsutella  Eutypa  Dothideales Pyrenophora Aspergillus Rhodotorula  Pseudohyphozyma  Dioszegia  Coprinopsis Pleurotus  Malassezia  Malassezia  Ceraceosorus Tilletia  Tilletia |
| --- | --- | --- |
| Claster 6 | ABK76739.1 | Cenarchaeum |

ADH87030.1

AEA60295.1

AEK46982.1

AEL11490.1

AFJ37304.1

AGM07905.1

AHF75075.1

ATX77085.1

AXA37448.1

AXG70223.1

BAB98371.1

BAG28863.1

BEK67628.1

CAA6800162.1 CAB39589.1

CAH1092428.1 CAH7666574.1 CAI2934356.1	CAI8025105.1	CAI8320368.1	CAL6010981.1 CCT60915.1

CDI53613.1	CRL43668.1 CRX63995.1 CVY03029.1 EAQ40622.1 EDM80849.1 EDY37525.1 EDY37601.1 EDZ42196.1 EDZ44726.1 EDZ62619.1 EEA92994.1 EEA93293.1 EEA95828.1 EEA96716.1 EEB78973.1 EEC18884.1 EED26966.1 EED31880.1 EED38259.1 EED40326.1

Desulfurivibrio Burkholderia

Amycolatopsis Streptococcus	Mycobacterium Amycolatopsis Candidatus

Reinekea

Candidatus Kordia

Corynebacterium Kocuria

Kitasatospora

uncultured

Mycobacterium Acinetobacter

Phakopsora	Aminobacter Geodia

Acidimicrobiaceae Hexamita

Acetobacter

Melanopsichium Sodalis

Acinetobacter

Staphylococcus Polaribacter

Plesiocystis Cyanobium Cyanobium

Rhodobacteraceae Rhodobacterales

Sulfurimonas Pseudovibrio Pseudovibrio Pseudovibrio Pseudovibrio marine

Ixodes Vibrio gamma

Stenotrophomonas Stenotrophomonas

EEE38053.1

EEE38159.1

EGF92247.1

EPB77601.1

GAA1250752.1 GAA1532396.1 GAA3718174.1 GAA4221180.1 GAL34627.1

GEA54767.1

GEK15721.1

GGL79115.1

GGS93162.1

GHE75466.1

GHJ37203.1

GLZ80466.1

KAE8820852.1 KAF0817652.1 KAF0852878.1 KAF8749932.1 KAH7815917.1 KAI8202438.1	KAI8261190.1	KAJ4474736.1 KAJ6463181.1 KAJ7314916.1 KAK1981815.1 KAK4159053.1 KAK6211714.1 KHJ95189.1

KHO51823.1 KHO54368.1 KIE02908.1

KJH45734.1	KJX94738.1	KKQ17675.1 KKR78658.1 KKW19200.1 KKW46762.1 KKY16046.1 KLU59756.1 KNG51820.1 KPA41092.1

Rhodobacteraceae Rhodobacteraceae Asticcacaulis

Ancylostoma

Kitasatospora	Brevibacterium Nonomuraea

Streptosporangium

Vibrio

Burkholderia Aliivibrio

Streptomyces Streptomyces Streptomyces Streptomyces

Actinorhabdospora

Hordeum

Bacillus

Andalucia

Rhizoctonia

Monocercomonoides

Colletotrichum Colletotrichum Lentinula

Mycena

Mycena

Colletotrichum

Cladorrhinum

Colletotrichum

Oesophagostomum archaeon

archaeon

Metarhizium

Dictyocaulus

Zymoseptoria Candidatus

Candidatus

Parcubacteria

Parcubacteria

Phaeomoniella	Peptococcaceae Stemphylium

Fusarium

KPU42524.1 LAF73089.1 OAP20665.1 OLN97762.1 OLQ09510.1 PCI24288.1

PFN43704.1	PMB77730.1 PRX05621.1	PVU29550.1 QDV92069.1 QES32490.1	QHO39439.1 QRV88233.1 QRW03712.1 QRW08027.1 QRW08254.1 RAV32734.1 RCN32765.1 RED44201.1 RUP91937.1	SCC38059.1	SDD57756.1 SDM13712.1 SDW88478.1 SEC02485.1	SEW34912.1 SFN35979.1	SFO61932.1	SKP74957.1	SLN03961.1	SMQ18894.1 SPE22833.1	STY09675.1	STY84553.1	STZ44403.1	SUV64836.1 TCK94635.1 TDP71971.1	TNC86776.1 TQL74901.1	TQN75117.1 TVY93831.1

Oxobacter

Aduncisulcus	Amycolatopsis Colletotrichum Symbiodinium SAR324

Bacillus

Fervidicoccus Actinomadura

Yersinia

Phycisphaerae Streptomyces

Arachis

Ceratobasidium	Ceratobasidium	Ceratobasidium	Ceratobasidium	Corynebacterium Ancylostoma

Aestuariispira

Corynebacterium Bacillus

Streptomyces

Halarsenatibacter Myxococcus

Arthrobacter

Austwickia

Xenorhabdus

Amycolatopsis

Mycobacteroides Corynebacterium Streptomyces

Candidatus

Legionella	Mobiluncus

Mycolicibacterium Bordetella

Paraburkholderia Bradymonas

Alcanivorax

Stackebrandtia Colletotrichum Lachnellula

TWB86807.1 USP77263.1	VFJ66424.1

WP_030212554.1 WP_073717513.1 WP_082092581.1 WP_086489236.1 WP_091419189.1 WP_093509194.1 WP_093809521.1 WP_ 111935001.1 WP_ 138465607.1 WP_ 138694967.1 WP_ 158774678.1 WP_ 166635816.1 WP_ 199892900.1 WP_201953563.1 WP_210841406.1 WP_221356924.1 WP_221586502.1 WP_234828179.1 WP_235018218.1 WP_247406099.1 WP_250225622.1 WP_255949096.1 WP_272135776.1 WP_275353334.1 WP_282381768.1 WP_284317101.1 WP_290199153.1 WP_303570735.1 WP_308151634.1 WP_319931012.1 WP_320411520.1 WP_326442843.1 WP_336697731.1 WP_342709929.1 WP_352757507.1 WP_355528695.1 WP_371314008.1 XP_004345918.1 XP_009266725.1 XP_014654155.1

Bradyrhizobium Curvularia

Candidatus

Streptomyces Bowdeniella	Devosia

Thioflexithrix

Micromonospora Sphingopyxis

Stappia

Paraburkholderia Poseidonocella

Nonomuraea

Cobetia

Idiomarina

Streptomyces Nocardia

Nocardiopsis	Streptomyces	Qipengyuania Sinorhizobium

Thermomonospora

unclassified

Chryseobacterium Brucepastera

Stigmatella

Xenorhabdus Arthrobacter

Labrys Zwartia Cobetia

Clostridium

Xenorhabdus

Candidatus

Streptomyces

Curtobacterium Bradyrhizobium Mesorhizobium Embleya

Hyphomonas	Acanthamoeba Wallemia

Moesziomyces

Note: 6 representative enzymes are shown with underline.

Table S3 Kinetic assays of cysteine synthases with different substrates.

|  |  | *km* (μM) | *V_max_*  (μM·min^-1^·mg^-1^) | *Kcat* (S^-1^) | *Kcat*/*Km* (S^-1^M^-1^) |
| --- | --- | --- | --- | --- | --- |
| RpCysK | NaHS | 3192.5±892.5 | 0.581±0.086 | 13.833±2.048 | 4.333×10^3^ |
|  | GSSH | 202.75±71.75 | 0.280±0.026 | 6.667±0.619 | 3.300×10^4^ |
|  | Na2S2O3 | 1205.9±456.1 | 0.021±0.003 | 0.495±0.076 | 0.410×10^3^ |
| RpCysM | NaHS | 3116.5±780.5 | 0.572±0.075 | 13.059±1.713 | 4.190×10^3^ |
|  | GSSH | 192.75±61.25 | 0.258±0.0196 | 5.890±0.448 | 3.068×10^4^ |
|  | Na2S2O3 | 1998.5±733.5 | 0.026±0.004 | 0.594±0.091 | 0.297×10^3^ |
| EcCysK | NaHS | 389±43 | 0.139±0.031 | 4.635±1.039 | 1.033×10^4^ |
|  | GSSH | 265±37 | 0.163±0.019 | 9.778±0.665 | 2.406×10^4^ |
|  | Na2S2O3 | 1036±82 | 0.074±0.028 | 2.454±0.935 | 1.731×10^3^ |
| EcCysM | NaHS | 267±54 | 0.143±0.019 | 4.79±0.63 | 1.956×10^4^ |
|  | GSSH | 325±33 | 0.155±0.046 | 5.16±1.55 | 1.467×10^4^ |
|  | Na2S2O3 | 408±28 | 0.173±0.028 | 5.81±0.92 | 1.262×10^4^ |
| SiRe_ 1641 | NaHS | 157±14 | 0.259±0.061 | 8.661±2.015 | 6.418×10^4^ |
|  | GSSH | 279±32 | 0.174±0.038 | 4.919±1.266 | 1.365×10^4^ |
|  | Na2S2O3 | 1126±162 | 0.065±0.011 | 1.521±0.511 | 1.031×10^3^ |
| Cys11 | NaHS | 849±51 | 0.186±0.013 | 3.127±0.208 | 4.208×10^3^ |
|  | GSSH | 426±34 | 0.125±0.064 | 2.083±0.335 | 5.823×10^3^ |
|  | Na2S2O3 | 3774±353 | 0.064±0.008 | 1.067±0.126 | 0.281×10^3^ |

Strain or plasmid	Characteristic or target protein	Source

Strains

| *E. coli* MG1655 | Used for amplied the EcCysM, EcCysK, CysJ and CysI fragments | Our lab |
| --- | --- | --- |
| *R. pomeroyi* DSS-3 | wide type, used to amplied the RpCysM and RpCysK fragments | a gift from prof. Yuzhong Zhang |
| *R. pomeroyi* DSS-3 ∆*sqrΔfccAB* | *sqr,fccA* and *fccB* deleted | Our lab |
| *S. islandicus* REY15A | Used to amplied the SiRe_ 1641 fragments | a gift from prof. Yulong Shen |
| *S. cerevisiae* S288C | Used to amplied the MET5 and MET10 fragments | Our lab |
| *S. pombe* YHL6381 | Used to amplied the Sir1 fragments | a gift from prof. Huang Ying |
| *E. coli* XL1-Blue MRF’ | Used for plasmid construction | Our lab |
| *E.coli* BL21(DE3) | Used for protein expression | Our lab |
| Plasmids |  |  |
| pET30a-EcCysM | Used to express the EcCysM protein | study  This |
| pET30a-EcCysK | Used to express the EcCysK protein | study  This |
| pET28a-RpCysM | Used to express the RpCysM protein | study  This |
| pET28a-RpCysK | Used to express the RpCysK protein | study  This |
| pET30a-SiRe_ 1641 | Used to express the SiRe_ 1641 protein | study  This |
| pET28a-CysJ | Used to express the CysJ protein | study  This |
| pET28a-CysI | Used to express the CysI protein | study  This |
| pET28a-MET5 | Used to express the MET5 protein | study  This |
| pET28a-MET10 | Used to express the MET10 protein | study  This |
| pET28a-SPO2634 | Used to express the SPO2634 protein | study  This |
| pET28a-Sir1 | Used to express the Sir1 protein | study  This |
| pBBR1MCS5 | Used to as contrast | study  This |
| pBBR1MCS5-CysJI | Used to construct CysJI overexpression strains | study  This |
| pBBR1MCS5- Met10&Met5 | Used to construct Met5&Met10 overexpression  strains | study  This |
| pBBR1MCS5-Sir1 | Used to construct Sir1 overexpression strains | study  This |
| pBBR1MCS5-SPO2634 | Used to construct SPO2634 overexpression  strains | study  This |
| pJFE3 | Used to as contrast | study  This |
| pJFE3-MET5&MET10 | Used to construct MET5& MET10 overexpression strains | study  This |

Table S5. Primers used in this study

| Primer name | Sequence (5'-3') | Descriptions |
| --- | --- | --- |

pET30a-F

pET30a-R

EcCysM-pET30a-F

EcCysM-pET30a-R

EcCysK-pET30a-F

EcCysK-pET30a-R

SiRe_ 1641-pET30a-F SiRe_ 1641-pET30a-R

pET30a-YZ-F

pET30a-YZ-R

pET28a-RpCysM-F pET28a-RpCysM-R RpCysM-pET28a-F RpCysM-pET28a-R pET28a-RpCysK-F pET28a-RpCysK-R RpCysK-pET28a-F RpCysK-pET28a-R pET28a-CysJ-F

pET28a-CysJ-R

CysJ-pET28a-F

CysJ-pET28a-R

pET28a-cysI-F

pET28a-cysI-R

cysI-pET28a-F

cysI-pET28a-R

pET28a-MET5-F

pET28a-MET5-R

MET5-pET28a-F

MET5-pET28a-R

pET28a-MET10-F

pET28a-MET10-R MET10-pET28a-F

MET10-pET28a-R pET28a-SPO2634-F pET28a-SPO2634-R SPO2634-pET28a-F SPO2634-pET28a-R pET28a-sir1-F

pET28a-Sir1-R Sir1-pET28a-F

GCACTCGAGCACCACCACCAC

CATATGTATATCTCCTTCTTAAAG

TTAAGAAGGAGATATACATAGTGAGTACATTAGAACAAACAATAG TGGTGGTGGTGGTGCTCGAGAATCCCCGCCCCCT

TTAAGAAGGAGATATACATAATGAGTAAGATTTTTGAAGATAACT TGGTGGTGGTGGTGCTCGAGCTGTTGCAATTCTTTCTC

TTAAGAAGGAGATATACATATGGTGTCAAACCCTCTTCACGTATTC TGGTGGTGGTGGTGCTCGACAGTTATTTGCTTACTCCTAAATCTCTT TGCTAGTTATTGCTCAGCGGT

AGCCCAGTAGTAGGTTGAGG

GGACGTCTGAAAGCTTGCGGCCGCACTCGA TGCGCATATGGCTGCCGCGCGGCAC

CGCGGCAGCCATATGCGCATTGCACGCGATC

CCGCAAGCTTTCAGACGTCCTCGAACACACCCG TATCAGTACTGAAAGCTTGCGGCCGCACT

GATCATATGGCTGCCGCGCGGCAC

GCGCGGCAGCCATATGATCTTAACCACCAC	CAAGCTTTCAGTACTGATATCCGGGAGTCGA

GTCTACTAAAAGCTTGCGGCCGCACTC GTCGTCATATGGCTGCCGCGCGGCAC

GCGGCAGCCATATGACGACACAGGTCC GCCGCAAGCTTTTAGTAGACATCTCGCT GGGATTAAAAGCTTGCGGCCGCACT

GCTCATATGGCTGCCGCGCGGCAC

GCGCGGCAGCCATATGAGCGAAAAACA

GCCGCAAGCTTTTAATCCCACAAATCACG TGCCTATTAAAAGCTTGCGGCCGCACTC

GTCATATGGCTGCCGCGCGGCAC

GCGCGGCAGCCATATGACTGCTTCTGACC

CGCAAGCTTTTAATAGGCATCTTCAGACACATCTT AAGCTTGCGGCCGCACTCGA

GGCATATGGCTGCCGCGCGGCAC

CGCGCGGCAGCCATATGCCAGTTGAGTTTG

CGAGTGCGGCCGCAAGCTTTTAGTAGACTTCTAAAA GCCGCTTGAAAGCTTGCGGCCGCACT

ATACATATGGCTGCCGCGCGGCAC

GCGCGGCAGCCATATGTATCGCTATTCCGA GCCGCAAGCTTTCAAGCGGCATCGGCGC

TAGAAGCTTGCGGCCGCACTCGAG TCATATGGCTGCCGCGCGGCAC

CCGCGCGGCAGCCATATGAGTGTAAAGGCTA

Primers for cloning the pET30a vector

EcCysM expression

EcCysK expression

SiRe_ 1641 expression

Primers for sequencing

RpCysM expression

RpCysK expression

CysJ expression

CysI expression

MET5 expression

MET10 expression

SPO2634 expression

Sir1 expression

Sir1-pET28a-R pET28a-YZ-F	pET28a-YZ-R

pBBR1MCS5-F

pBBR1MCS5-R

CysJ-pBBR1MCS5-F CysI-pBBR1MCS5-R SPO2634-fragment-F SPO2634-fragment-R pBBBR5-YZ-F

pBBBR5-YZ-R	pJFE3-vector-F	pJFE3-vector-R	pJFE3-MET5-F	pJFE3-MET5-R	pJFE3-MET10-F pJFE3-MET10-R pJFE3-YZ-F

pJFE3-YZ-R

GTGCGGCCGCAAGCTTCTAAAACGCTTCAGGT

GCTCATGAGCCCGAAGTGGC

GCGAGAAAGGAAGGGAAGAAAGCG	TGAGTCGTATTACGCGCGCTCACTGG TAGGGCGAATTGGAGCTCCACCGC

GGAGCTCCAATTCGCCCTAATGACGACACAGGTCC

GCGCGCGTAATACGACTCATTAATCCCACAAATCACG GGAGCTCCAATTCGCCCTAATGTATCGCTATTCCGAAT GCGCGCGTAATACGACTCATCAAGCGGCATCGGCGC

GATTCATTAATGCAGCTGGCAC

GTTTAAGGGCACCAATAACTGCC

AGTCGACCTGCAGGATTGAATTGAATTGAAAT

TAGCAAACTCAACTGGCATCTAGAGGATCCTTTGTAATTAAAAC GGATCCTCTAGATGACTGCTTCTGACCTCT

TCAATCCTGCAGGTCGACTTTAATAGGCATCTTCAGAC ATGCCAGTTGAGTTTGCTACCAATCCTTT

GCAGTCATCTAGAGGATCCTTAGTAGACTTCTAAAATGTATCT GTGATCCCCCACACACC

GACCATGATTACGCCAAGC

Primers for sequencing

Primers for cloning the pBBR1MCS5 vector

Primers for cloning the *cysJI* fragments

Primers for cloning the *spo2634* fragments

Primers for sequencing

Primers for cloning the pJFE3 vector

Primers for cloning the *met5* fragments

Primers for cloning the *met10* fragments

Primers for sequencing

Underlining represents the sequences of homologous arms.
